# Spanning spatial scales with functional imaging in the human brain at 10.5 Tesla

**DOI:** 10.1101/2024.12.20.629800

**Authors:** Luca Vizioli, Lasse Knudsen, Yulia Lazarova, Steen Moeller, Logan Dowdle, Andrea Grant, Alireza Sadeghi-Tarakameh, Nils Nothnagel, Yigitcan Eryaman, Matt Waks, Russel L. Lagore, Edward Auerbach, Gregor Adriany, Federico De Martino, Lonike Faes, Essa Yacoub, Kamil Uğurbil

## Abstract

One of the most important frontiers in the effort to improve functional imaging of brain activity (fMRI) is the recent push to increase the magnetic fields for human imaging beyond 10 Tesla. Having established safety, large gains in signal-to-noise ratio (SNR), and novel radiofrequency arrays that capture the higher SNR, here we demonstrate major gains in the spatial resolution, sensitivity, and functional contrast of blood-oxygenation-level-dependent (BOLD) based human fMRI at 10.5 Tesla. We demonstrate that BOLD contrast increases supralinearly with magnetic field in going from 7 to 10.5T, and ultrahigh resolutions achieved at 10.5 Tesla suppress large-vein confounds in gradient-recalled-echo BOLD fMRI, thus improving the fidelity of functional signals relative to neuronal activity and yielding accurate cortical depth profiles for layer-specific activation. Together, these multiplicative benefits deliver much needed increases in precision and resolution for meso-scale fMRI applications and illustrate the transformative potential of human functional imaging at magnetic fields that exceed 10 Tesla.

## 1. INTRODUCTION

Following its introduction^1, 2^, functional MRI (fMRI) based on the blood-oxygen-level-dependent (BOLD) contrast has rapidly emerged as the most widely used noninvasive tool employed for human brain studies. Nevertheless, like all techniques at our disposal for studying the brain, fMRI has limitations, particularly with respect to spatiotemporal resolution and the fidelity of the functional maps relative to the neuronal activity that induces them.

The brain operates over a vast spatial scale, going from individual cells to mesoscale neuronal ensembles and ultimately to brain-wide networks involving billions of neurons. However, the vast majority of human fMRI experiments are performed with spatial resolutions that correspond to voxel volumes of ≳ 8 µL (e.g. ≥ 2 mm isotropic resolution), containing more than a million neurons. Despite the relatively high signal-to-noise ratio (SNR) of MR images at such low resolutions, detecting the inherently small functional signals remains challenging due to the low functional contrast of the fMRI method. Addressing this challenge often requires group-level studies involving a large number of subjects, and/or prolonged and repeated scanning sessions with single individuals (e.g.^3, 4^ and references therein). Aiming for higher spatiotemporal resolutions significantly exacerbates the problem due to the consequent decreases in SNR and functional contrast-to-noise ratio (fCNR).

An additional complication arises from the indirect nature of functional signals in fMRI, which reflect neuronal activity through neurovascular coupling. A well-known deleterious consequence of this indirect process is the draining-vein confound associated with gradient-recalled echo (GRE) BOLD fMRI (e.g.^5–12^), by far the most commonly employed fMRI technique for studying the brain. This confound leads to displacement and blurring of the recorded functional maps relative to the neuronal activity that elicits them.

Overcoming the afore-listed limitations has been a major research focus since the introduction of the fMRI technique; in this effort, the development of ultrahigh magnetic fields (UHF, defined as ≥ 7 Tesla) for human imaging (e.g., reviews^13–16^) has been one of, if not the most, consequential initiatives. Higher magnetic fields provide increases in SNR^17–21^, the magnitude of the BOLD effect (e.g. ^22–25^), and the relative contribution of the weak but spatially accurate functional signals associated with microvasculature (e.g., ^22, 23, 26^). With these advantages, 7 Tesla (7T) fMRI ushered in the ability to break the millimeter-resolution barrier and access mesoscale neuronal organizations (e.g., reviews ^8, 15, 16, 27–36^), starting with the demonstration of functional mapping of ocular dominance columns and orientation domains in the same individual^37^. In addition, 7T was shown to provide higher detection sensitivity and model predictive power (e.g.,^38–40^) at any spatial resolution relative to 3T.

Yet, reliably accessing mesoscale computations in the human brain remains challenging. A typical voxel size of ∼0.8 mm isotropic (∼0.5 µL voxel volume), accessible robustly and without excessive acquisition times at 7T, still contains more than forty thousand neurons^41^ capable of performing a multitude of different computations (e.g.,^42, 43^). This limitation was recognized in the strategic plan developed for the US BRAIN Initiative, which codified resolution targets of ≤ 0.1µL voxel volume initially ^44, 45^ and 0.01µL five years later^46^, emphasizing the *“ambitious*” and *“elusive*” nature of these targets.

One of the recent major initiatives undertaken to meet these challenges is the effort to push human imaging beyond 10T^47, 48^, spearheaded by the 10.5T effort in our laboratory^18–21, 49–53^. Having established safety^52, 54^, large SNR gains for human brain imaging^18–21^, and novel radiofrequency arrays that capture these SNR gains^20, 21^, we report the first results pushing the limits of functional imaging at this unique magnetic field, demonstrating major gains in spatial resolution and precision.

## 2. RESULTS

All resolutions given are isotropic and, unless otherwise specified, “nominal”. The *effective*voxel dimensions typically exceed the nominal resolution but are rarely reported. In this paper, when relevant, we report estimates of functional blurring (see Methods), which provide an indication of the *effective* resolution.

### 2.1. Quantitative T2* measurements

T2* is a critical parameter in fMRI (e.g.^5, 55^) and depends on the spatial resolution and the magnetic field used. Hence, measurements of T2* in the entire brain were performed at two different spatial resolutions and two different magnetic fields. We used a multi-echo gradient-recalled echo (MEGRE) sequence in the same subject at 10.5T to obtain whole-brain T2* measurements at 0.37- and 0.8-mm resolutions. A partial field of view acquisition (axial slab, centered around the calcarine fissure) at 0.35mm resolution was also obtained at 7T. Following processing and tissue segmentation, we obtained voxel-wise estimates of the T2* using a nonlinear fitting approach. For visualization of the full extent of data, we show an unweighted average of the 10.5T 0.37 mm data (Fig. 1a).

**Fig. 1:**
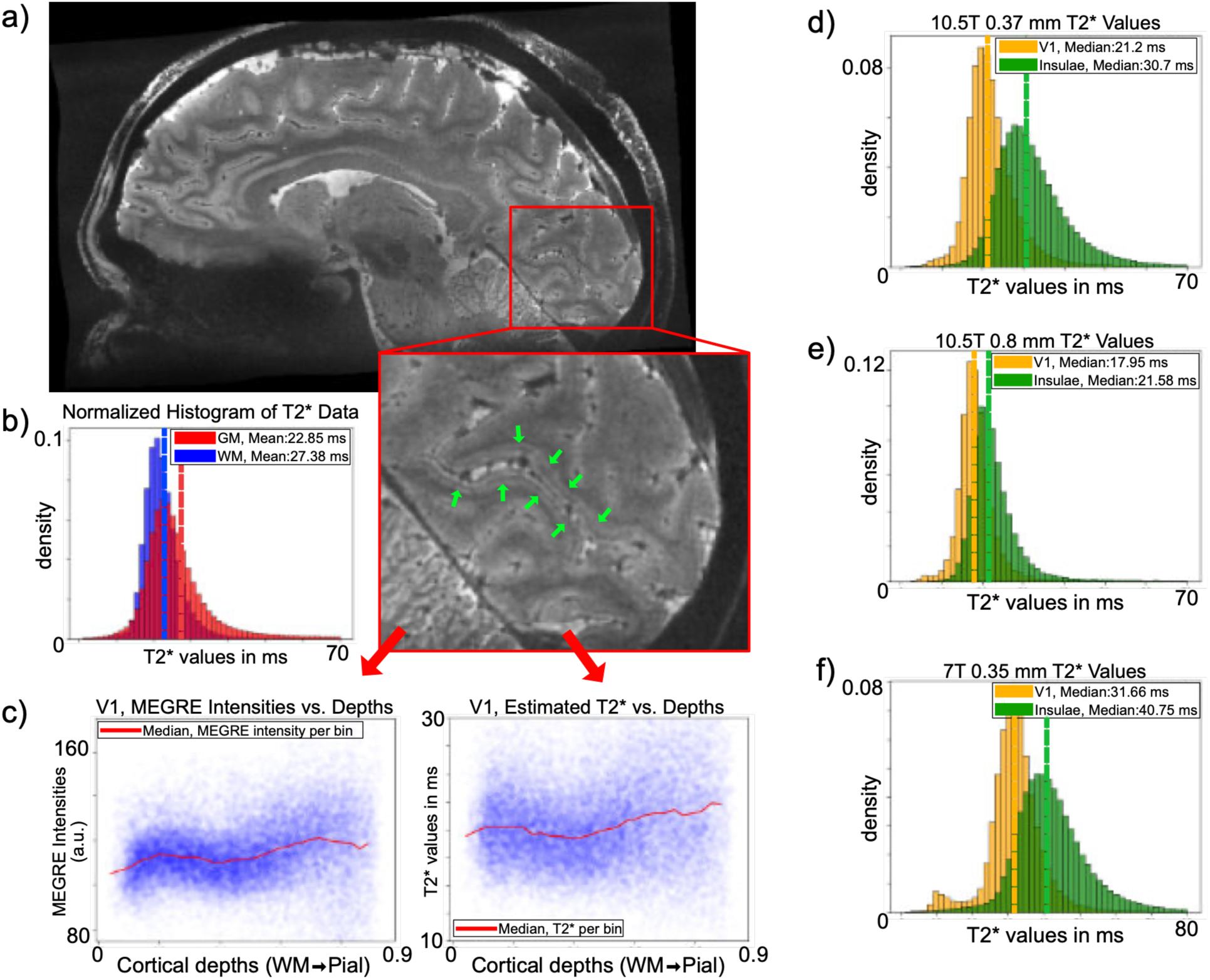
Quantitative T2* measurements at 10.5T obtained with MEGRE with 0.37 mm (2 runs, 7 echoes covering 4.07 – 27.4 ms) and 0.8 mm (3 runs, 7 echoes, 4 – 28 ms) resolutions (panels **a** through **e**) and 7T with 0.35 mm resolution (6 runs, 6 echoes, 4.37 – 26.22 ms) (panel **f**). **a**) (unweighted) average of all echoes from MEGRE runs at 10.5T with a 0.37mm isotropic spatial resolution, zooming in on the calcarine sulcus, where the Stria of Gennari (the dark thin line indicated by the green arrows), which runs in the middle of the cortex on both the upper and lower banks of the calcarine sulcus, is clearly visible. In these images, the cortex appears lighter gray compared to the white matter, and the myelinated fibers of the Stria of Gennari form a darker line in the middle of the cortex. **b)** Normalized histogram of T2* values (ms) for gray matter (red) and white matter (blue) for the 10.5T 0.37mm data. Dotted vertical lines represent the means. **c)** MEGRE intensities (T2* weighted average, restricted to V1, out to an eccentricity of 90 degrees, left panel) and estimated T2* values (ms) across cortical depths. Panels **d-f** show the histograms of T2* values (ms) for V1 (yellow) and insulae (green) for 10.5T 0.37mm data **(d)** and 0.8mm data **(e)**, and for 7T 0.35mm data **(f).** Dotted lines represent the median.

The distributions of gray and white matter T2* values (Fig. 1b) were derived from tissue segmentations in the 10.5T 0.37mm data and show high variability across the cortex. Using the gray matter segmentation, we examined the T2* as a function of depth within a region in the primary visual cortex (V1). We observe a decrease in T2* (Fig. 1a, expanded panel, and Fig. 1c) at the mid-cortical depth, expected due to the presence of myelinated fibers associated with the Stria of Gennari in the middle layer of V1. This effect was also visible in the signal intensity of the weighted average images (Fig. 1c, left panel).

T2* was further investigated in the primary visual cortex and the insulae, by combining gray matter segmentations with atlas-derived measures of these regions (see Methods). The 10.5T 0.37 mm data (Fig. 1d) showed a marked increase in T2* estimates relative to the 0.8mm data at the same field strength (Fig. 1e) for both regions. In the visual cortex, gray matter T2* (median) at 10.5T was 21 ms vs. 18 ms for 0.37- and 0.8-mm resolution data, respectively. For the insulae at 10.5T, the corresponding numbers were 31 ms vs. 22 ms, respectively. The T2* estimates with 0.8 mm resolution at 10.5T were similar to what is expected based on extrapolations from 9.4T data^17^. At 7T, 0.35mm acquisition yielded approximately 10 ms longer T2* for both the visual cortex and insulae gray matter relative to their 10.5T counterparts at a similarly high resolution (Fig. 1f), displaying the expected linear dependence of 1/T2* with the field magnitude (e.g. ^23^).

### 2.2. BOLD effect magnitude at 0.4 mm isotropic resolution: 7T vs 10.5T

We recorded 0.4 mm (i.e. 0.064 µL voxel volume) resolution functional images in three participants at 10.5T and 7T (Fig. 2) using otherwise matched approaches and parameters. Acquisitions were based on 3D GRE-EPI in a slab placed over the occipital cortex. The TE was ∼24 ms for both field strengths, which is near optimal for BOLD-fMRI at both fields, positioning the acquisition close to but on two different sides of the broad maximum defined by TE = T2* ^5, 55^. The experimental paradigm was that implemented in^56^, consisting of a 24-second on/off block design with a center and surround flickering checkerboard stimulation, and data were acquired in consecutive ∼5-min runs. Data were processed with the vendor default reconstruction *without* NORDIC denoising^56^.

**Fig. 2:**
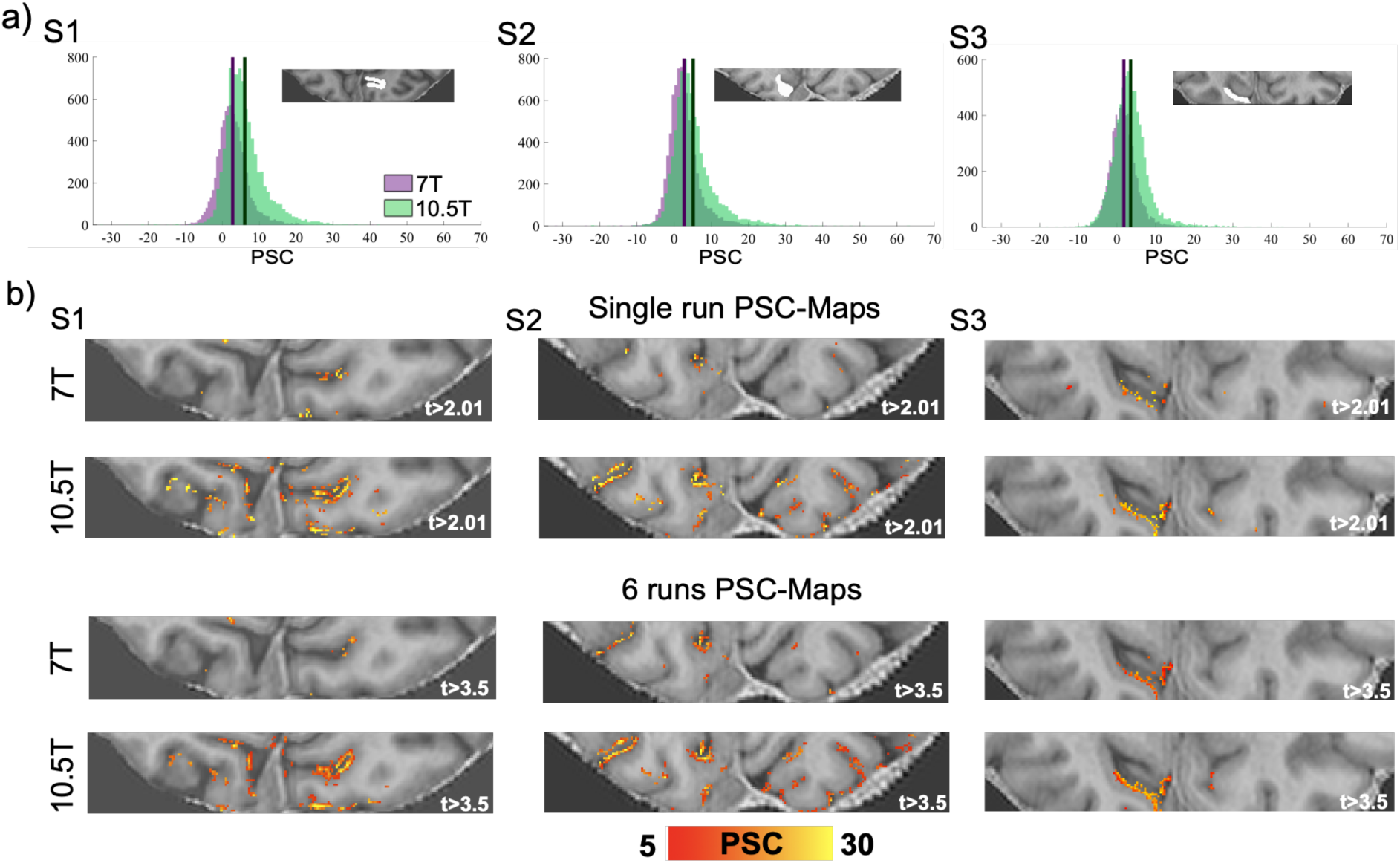
0.4 mm isotropic functional imaging of retinotopically organized visual stimulation at 7T and 10.5T obtained with parameters iPAT=3; PF=5/8; 40 Slices; TR=113 ms; VAT=4972 ms; TE=24.4 ms for 7T and 10.5T, with minor variations in TR=111 ms; VAT=4884 ms; TE=24.2 ms for the latter. **(a)** Histograms of voxel-wise PSC (computed across 5 runs - i.e. 25 minutes of data) for the contrast target and surround > 0 in 3 subjects. All voxels within the region of interest (example slice shown in the insert) were included. **(b)** 7T (top) and 10.5T (bottom) PSC maps superimposed on a co-registered MP2RAGE image for the contrast target and surround > 0, for a single run (top panel; t-threshold of 2.01, cluster threshold of 5) and for 6 concatenated runs (bottom panel; t-threshold of 3.5, cluster threshold of 5).

We carried out a region of interest (ROI) analysis (see Methods) confined to the retinotopic representation of target and surround in V1 (Fig. 2a). To avoid confounds related to SNR and BOLD response differences across field strengths, the ROI was anatomically defined using the Benson probabilistic atlas,^57^ guided by one functional run (which was excluded from further analyses - see Methods). Percent signal change (PSC) elicited by target and surround was estimated across the five remaining runs. The mean ratio between the PSC at 10.5T vs. 7T across participants (±STD) was 2.02 (±0.11) (PSC ratio for individual participants: S1 = 2.15; S2 = 1.93; S3 = 1.99). The mean PSC (±SEM) across all voxels within the ROI was 6.08±0.46 (S1), 5.11±0.34 (S2), and 3.52±0.21 (S3) at 10.5T and 2.83±0.29 (S1), 2.65±0.37 (S2), and 1.77±0.16 (S3) at 7T.

Fig. 2b shows activation maps as PSC for the same contrast target ∩ surround > 0 for a single run (top; t-threshold > 2.01) and for all 6 concatenated runs (bottom; t-threshold > 3.5) for both field strengths superimposed on representative slices of the co-registered MP2RAGE images.

For these three subjects, we computed the fCNR, as the mean BOLD PSC evoked amplitudes over the model residuals’ standard deviation. Without NORDIC denoising, the average fCNR for 7T was 0.12 (S1 = 0.13; S2 = 0.13; S3 = 0.1), while for 10.5T it was 0.24 (S1 = 0.24; S2 = 0.29; S3 = 0.18), representing a doubling of average fCNR at 10.5T relative to 7T, in this case reflecting almost exclusively the doubling in PSC. Since *fCNR* = (Δ*S*/*S*) · *tSNR*, where Δ*S*/*S* is PSC, these data indicate that with these acquisition parameters tSNR is similar for 7T and 10.5T data. However, these data were obtained with virtually the same TE (24.4ms (7T) and 24.2ms (10.5T)). Given the shorter T2* at 10.5T vs 7T, this penalizes the 10.5T tSNR.

In order to avoid the complications created by the presence of a task or stimulation, we recorded 2 additional human “resting state” (10 runs per participant) and 1 phantom tSNR (4 runs) data with identical acquisition parameters to the data in Figure 2. We observe a significant (p<0.05) albeit modest (when averaged over the whole slab) increase in tSNR at 10.5T vs 7T in the in vivo brain (∼20% see SUPPLEMENTARY Information, section 1). In contrast, in the phantom, tSNR was 2.75 times higher at 10.5T vs. 7T. (SUPPLEMENTARY Information, section 1).

### 2.3. 0.35 mm isotropic BOLD EPI images in visual cortex

Fig. 3 displays 0.35 mm (0.0428 µL voxel volume) resolution (3D GRE EPI) 10.5T images over the visual cortex for the same target-and-surround block design visual paradigm used in Fig. 2. Fig. 3a and the top panel in Fig. 3d show the t-maps for the target (in red) > surround (in blue) for an example subject for 6 consecutive 5-min functional runs. Functional maps are superimposed on slices from a single GRE EPI volume of the fMRI time series in panel a, and on an axial slice of the T_2_*-weighted images from the 0.37 mm resolution MEGRE acquisition in panel d. Single voxel timecourses for a single 5-min run for target-specific voxels are presented in Fig. 3b, clearly displaying remarkably robust signal intensity changes selectively associated with the target stimulus epochs (dark blue bands). These voxels were randomly selected amongst significantly active voxels within the retinotopic representation of the target in V1 (see Methods). Fig. 3c and 3d (bottom panel) show the t-maps (target > 0) evoked by single condition mapping for only 15 minutes of data in a different subject, superimposed on a selection of axial (top and middle panels in Fig. 3c) and coronal (bottom panel Fig. 3c) slices for the mean EPI, and on the MEGRE (Fig. 3d bottom). The green boxes in Fig. 3c zoom into cortical regions demonstrating the separation of activation across adjacent banks of a sulcus (blue arrows), regardless of the presence (leftmost panel) or absence (middle panel) of large vessels, for single condition GRE BOLD-fMRI. Moreover, as previously reported^58^, the green arrows in panel 3d indicate the Stria of Gennari, corresponding to layer 4B (Brodmann classification) in V1, clearly visible as a dark stripe traversing the cortical ribbon for both subjects.

**Fig. 3:**
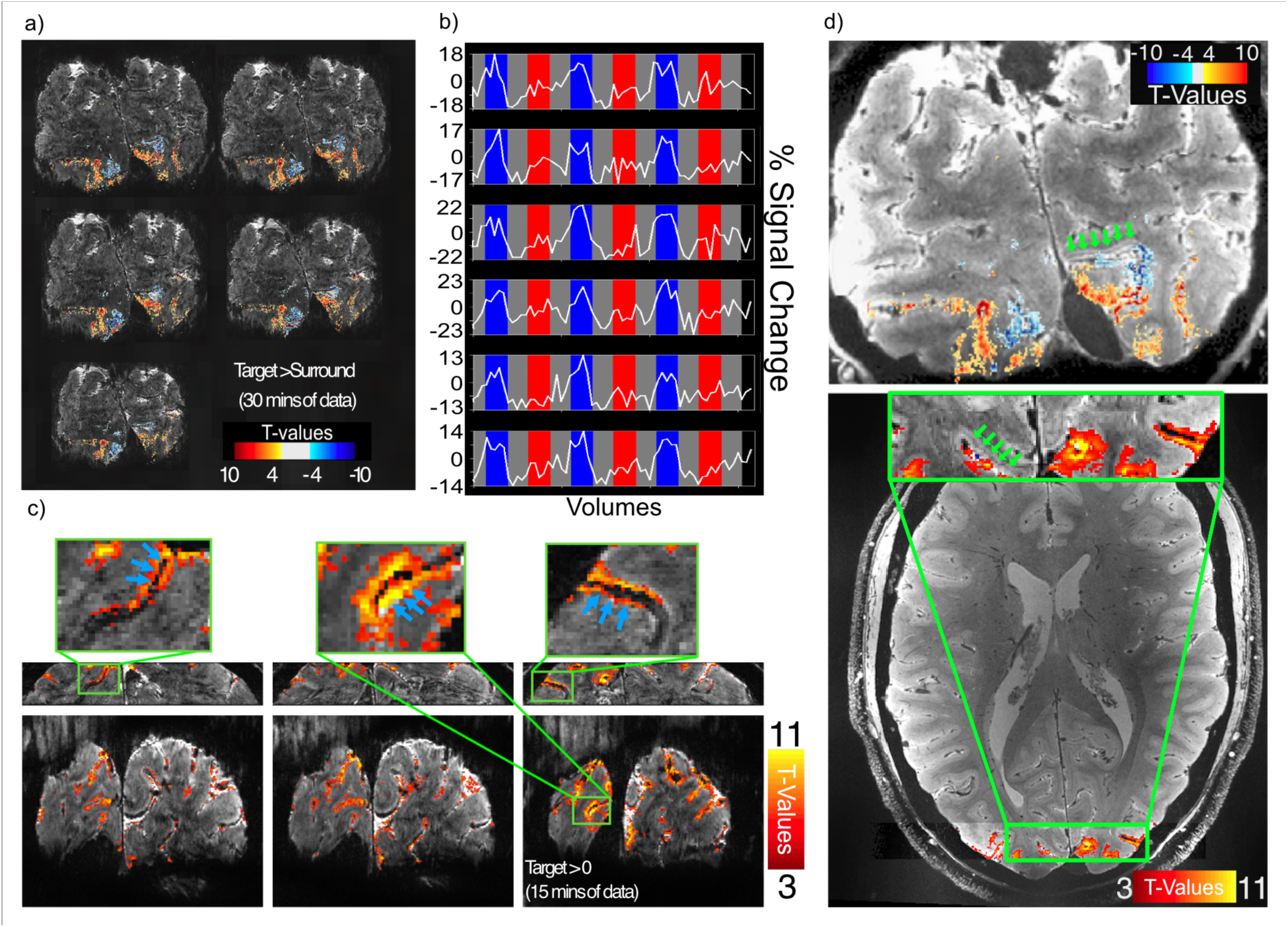
0.35 mm resolution functional images at 10.5T acquired with iPAT=4; 5/8 PF; 40 Slices; TR=113 ms; VAT=4972 ms; TE=24ms for 2 representative subjects. **(a)** t-maps for the contrast target > surround superimposed on single slice EPI images; **(b)** Examples of target-selective single-voxel single-run timecourses. The different colors represent different stimulation periods (dark blue: target; red: surround; gray: baseline); **(c)** t-maps for the contrast target > 0 superimposed on a selection of single slices mean EPI images; blue arrows on the zoom-in indicate how this resolution allows for the clear separation of the activation across adjacent banks of a sulcus, even for single condition maps, regardless of the presence (leftmost and rightmost panel) or absence (middle panels) of large veins **(d)** top panel: target > surround t-maps superimposed on a T2* image; bottom panel: target > 0 t-maps superimposed on a T2* image; green arrows indicate the Stria of Gennari (dark line). The artifact that appears in the upper left corner of Fig3,c, lower row of images comes from signals outside the field-of-view (FOV) aliasing into the image. With a coronal slice selection, one in principle gets signals from the brain and also from inferior regions such as the skull and even the neck. However, the FOV is set such that they lie outside the FOV but now fold (i.e.”alias”) into it, a common practice of letting such signals alias into a location where they do not overlap the signals of interest (in this case brain).

### 2.4. Layer profiles for 0.35 mm and 0.8 mm isotropic resolutions

Four subjects were scanned using the 0.35 mm functional acquisition and visual stimuli described in Fig. 3. The same four subjects were also scanned with 0.8 mm resolution at 7T (see Methods).

We first segmented the cortical ribbon into 13 depths within the retinotopic representation of the target stimulus in V1 with an equivolume approach (see Methods). The mean signal intensity of each depth for the 0.37 mm MEGRE images and the mean EPI images of the fMRI time series across both resolutions are plotted in Fig. 4a. The plots in Fig. 4a and 4b are color-coded so that all data are plotted in red for 10.5T and in blue for 7T. For all subjects, we observed a “*dip”* in the MEGRE data (obtained at 10.5T) at approximately 50% of the distance into the cortex from the cerebrospinal fluid (CSF) boundary (Fig. 4a). The same *“dip”* is clearly visible for the 10.5T 0.35 mm mean GRE EPI images of the fMRI time series (dashed red lines in Fig. 4a) but not for the 0.8 mm mean GRE EPI images of the 7T fMRI time series (blue dashed lines, Fig. 4a). Expanded versions of this figure are supplied in SUPPLEMENTARY figures. S9.1 and S9.2.

**Fig. 4:**
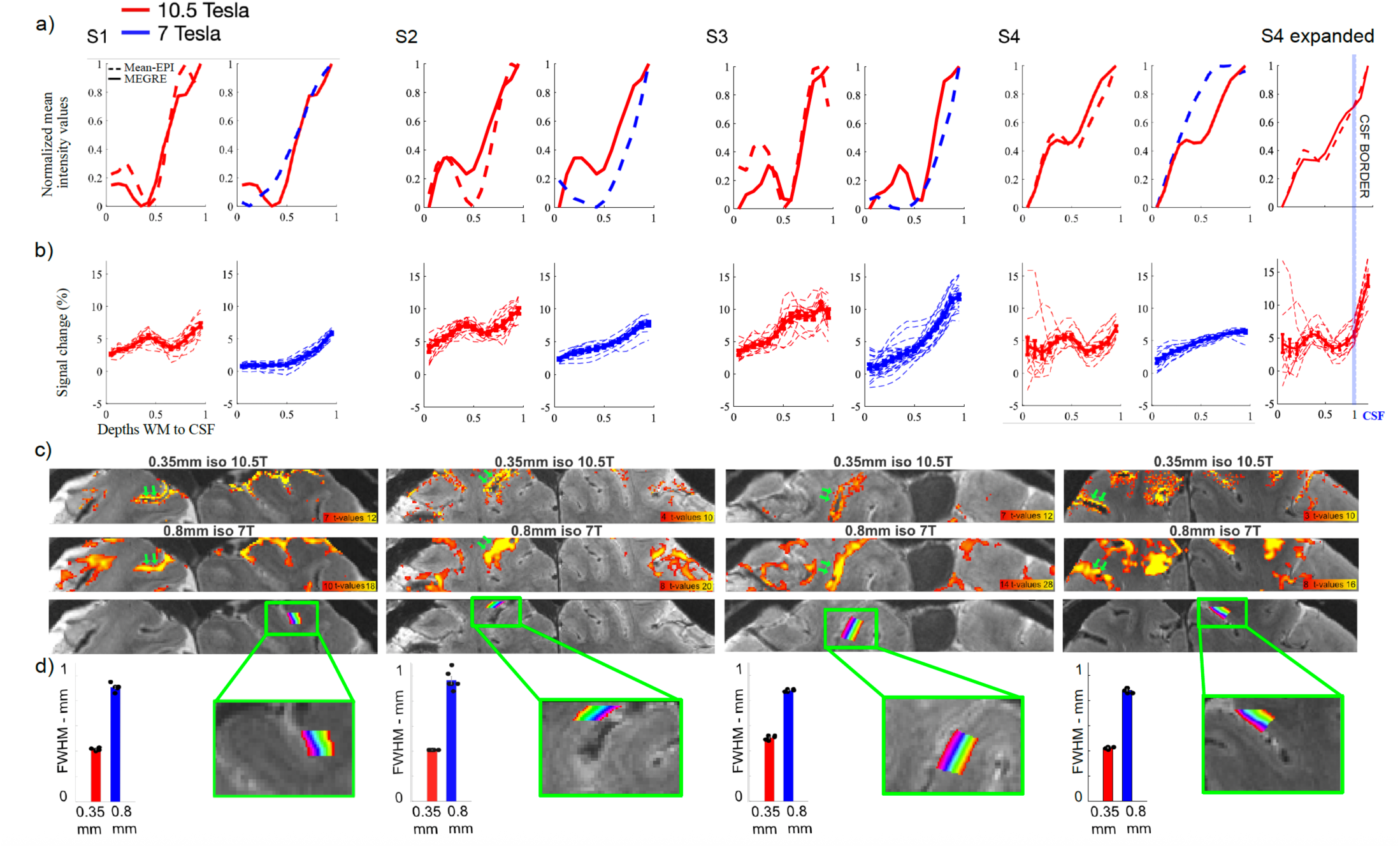
Comparison of 0.35 mm (10.5T) and 0.8 mm (7T) functional data. Panel (**a)** depicts laminar profiles of normalized (min=0, max=1) signal intensities for the T2*-weighted acquisitions (mean EPI and MEGRE images). The dashed lines refer to the mean EPI intensity profiles (red: 10.5T 0.35 mm; blue: 7T 0.8 mm). The MEGRE data are from 10.5T and have 0.37 mm isotropic resolution (solid red line, included in both panels of each subject). The dip in the middle depths seen in the 10.5T MEGRE and 0.35 mm mean EPI profiles identifies the location of the Stria of Gennari, which is not visible in the corresponding 0.8 mm 7T mean-EPI profiles. Panel **(b)** shows the functional PSC laminar profiles for the target condition. For all subjects, the 10.5T 0.35 mm PSC profiles (in red) demonstrate a bump co-localized with the dip seen in the corresponding mean-EPI and MEGRE anatomical traces in panel (a). This bump was not resolved at 0.8 mm resolution at 7T (blue). Error bars represent standard error of the mean (SEM) across single trials. **(b)** PSC in the 10.5T 0.35 mm data, showing a rapid increase immediately beyond the boundary between cortex and CSF (light-blue vertical line). **(c)** Illustration of the cortical layers used to extract laminar profiles in panel *a)* from an example slice in each subject (lower panel). The top and the middle panels show activation (T-maps, cluster threshold of 10) for the 0.35 mm and 0.8 mm acquisitions, respectively. The green arrows point to example locations where activation across adjacent sulcal banks is separable at 0.35 mm but not at 0.8 mm resolution. **(d)** Bar plots depict the spatial autocorrelation-based blurring estimates, showing significantly lower smoothness in the 0.35 mm data; error bars represent SEM across imaging runs.

We further extracted the average PSC from the fMRI data within each depth elicited in all individual ∼5-min runs (dashed lines in Fig. 4b). The average of these single-block PSC profiles is shown as a solid line in the same panels. Note that the single 5min runs display essentially the same profile with little scatter. The laminar data lead to three main observations:

The 0.35 mm 10.5T functional activation profiles (Fig. 4b) exhibited a local peak/bump co-localizing with the dip in the corresponding mean-EPI profile (Fig. 4a), which was significant across trials in all subjects (see Methods; S1: t(11)=3.68, p=0.002; S2: t(11)=6.30, p<0.001; S3: t(11)=2.34, p=0.020; S4: t(11)=3.24, p=0.004). This peak/bump was not observed in the 0.8 mm data (S1: t(11)=0.74, p=0.237; S2: t(11)=0.19, p=0.425; S3: t(23)=-0.14, p=0.557; S4: t(11)=0.195, p=0.425).

The average slope of the ramping profile displaying increasing PSC towards the pial surface (i.e. towards CSF) – typical of GRE BOLD fMRI - was significantly shallower for the 0.35 mm 10.5T fMRI data compared to the 0.8 mm 7T fMRI images, as evaluated with two-sample t-tests (S1: (mean±Std across trials) 0.35mm: 3.85±1.76, 0.8mm: 5.93±0.97, t(22)=-3.61, p=0.002; S2: 0.35 mm: 6.01±1.24, 0.8 mm: 7.47±2.03, t(22)=-2.12, p=0.045; S3: 0.35 mm: 8.12±2.09, 0.8 mm: 12.63±2.87, t(22)=-4.41, p<0.001; S4: 0.35 mm: 1.81±5.92, 0.8 mm: 5.75±2.06, t(22)=-2.18, p=0.040 – See SUPPLEMENTARY Fig. S8). This was quantified by fitting a line to the PSC laminar profile elicited by each block and estimating its slope (see Methods). This result was primarily driven both by the apparent increase in evoked PSC amplitudes in the inner depths and the decreased amplitudes towards the pial surface for the 10.5T 0.35 mm data, compared to the 7T 0.8 mm data.

Finally, in Fig. 4a and 4b, the rightmost column labeled “S4 expanded”, reproduces the 10.5T 0.35 mm data for subject S4, over an expanded depth, showing a rapid increase in PSC immediately beyond the GM-CSF boundary of the cortex.

The 0.35mm and 0.8mm resolution functional maps for each participant are shown in Fig. 4c. The 0.8mm activation maps are upsampled to the grid of the 0.35mm data (see SUPPLEMENTARY Information, Fig. S7 for 0.8 mm activation maps in their native resolution). As expected from a retinotopic paradigm such as the one employed here, the spatial extents and locations of activation look highly comparable across resolutions. However, the activation between adjacent banks of a sulcus (indicated by the green arrows in Fig. 4c (also see Fig. 3c)) are clearly separable for the 0.35 mm 10.5T images but, as expected, not for the 0.8 mm 7T ones, which are blurred into a single cluster.

The global smoothness of the fMRI time series within the visual cortex in EPI images was estimated using AFNI (3dFWHMx function)^59^ (Fig. 4d). Specifically, spatial autocorrelation was computed from the general linear model (GLM) residuals using a Gaussian + monoexponential decay mixed model to account for possible long-tail autocorrelations.

The functional resolution estimates (mean±Std across runs) were 0.41±0.01mm (S1), 0.41±0.00mm (S2), 0.50±0.02mm (S3), and 0.42±0.01mm (S4) for the 10.5T 0.35mm data and 0.91±0.03mm (S1), 0.97±0.07mm (S2), 0.88±0.01mm (S3), and 0.88±0.02mm (S4) for the 7T 0.8mm data. Two-sample t-tests carried out across runs independently per subject showed that the 10.5T 0.35mm data were significantly less smooth than the 0.8mm data (S1: t(10) =-39.55, p<0.001; S2: t(10) = −18.38, p<0.001, S3: t(10) =-49.55, p<0.001; S4: t(10) = −52.92, p<0.001).

### 2.5. 0.65 mm isotropic resolution GRE BOLD EPI images in auditory cortex at 10.5T

It is not possible to cover the whole brain or large sections of the brain without prolonged volume acquisition times (VATs) with very high resolutions, such as 0.4 mm and 0.35 mm isotropic. Here we demonstrate the quality of data obtained within reasonable VATs at a resolution that still exceeds what has been typically possible at lower magnetic fields and signal strengths. Fig. 5 displays functional images obtained without denoising, covering a large fraction of the brain during auditory stimulation at 0.65 mm isotropic resolution, the highest resolution used for the auditory cortex to date, using a mixed fast-event-related/block design auditory stimulation paradigm (see Methods).

**Fig. 5:**
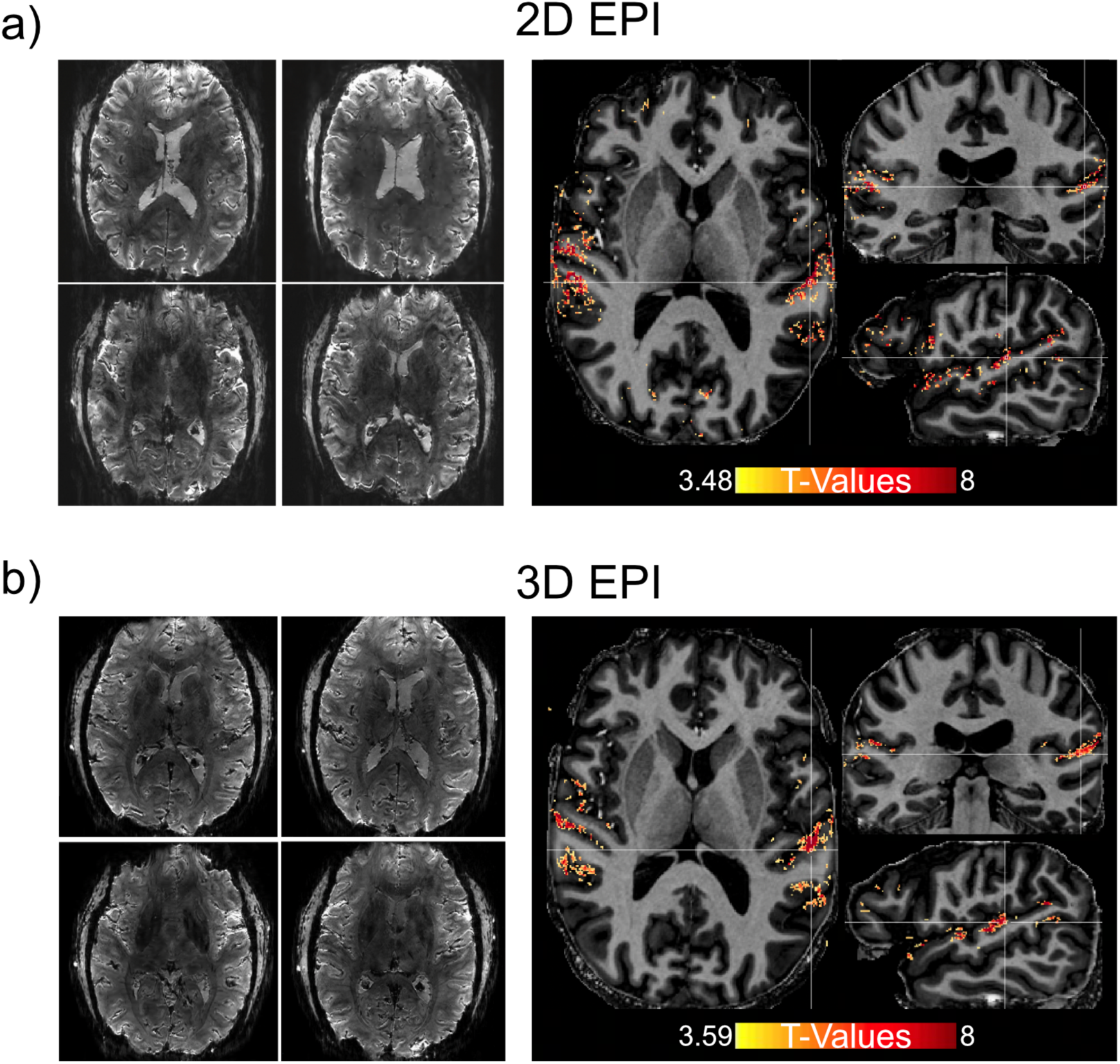
0.65 mm isotropic (nominal) resolution GRE BOLD fMRI with auditory stimulation for **(a)** 2D (TE=20.6 ms; TR=2320 ms; in-plane acceleration iPAT=4; MB=2; PF=5/8; 60 Slices; FOV=(296×292 mm)) and **(b)** 3D EPI (TE=16.6 ms; TR=81 ms; in-plane acceleration iPAT=4; PF=5/8; 52 Slices; VAT=4536 ms, FOV=(272×276 mm)). For both (a) and (b), the left panels show an example of single slices of a single EPI volume. The panel on the right shows activation to sounds (t-maps for sounds > 0) for 5 concatenated runs (just over 35 minutes of data) superimposed on a co-registered T1 (MP2RAGE) for the offline tailored reconstructions.

Functional maps were acquired with GRE EPI using Simultaneous Multi-Slice (SMS)/ Multiband (MB) ^60–62^ and 3D EPI strategies and are shown in Fig. 5 for t-threshold (t>3.5) and (t>3.6), respectively, and a cluster threshold of 4 voxels for the sounds > 0 contrast across the auditory cortex for 5 concatenated runs. Each run was approximately 8 min of data. VAT was 2.3s in SMS/MB and 4.5s in the 3D EPI. 5/8 partial Fourier was used to reduce the TE below optimal for BOLD contrast to minimize the volume acquisition times.

### 2.6. Whole-brain fMRI with 0.75 mm resolution BOLD EPI

Reducing the resolution to 0.75 mm isotropic (i.e. 0.42 µL), we were able to extend the coverage further relative to data presented in Fig.5 and cover the whole brain with a temporal resolution of ∼3.2 seconds. The quality of the data acquired at 10.5T with SMS/MB is illustrated in Fig. 6. The acquisition parameters were: 0.75mm isotropic voxels; 12-fold 2D acceleration (MB=3; in-plane (iPAT)=4); PF=5/8; 144 Slices; TR=3210 ms; TE=18 ms. We used a 24-second on/off block design visual paradigm. Image acquisition and reconstruction were performed using the HCP Multiband reconstruction developed in CMRR (downloadable from https://www.cmrr.umn.edu/multiband/), *without denoising* and with minimal preprocessing (see Methods). Fig. 6 shows the activation maps elicited by faces, noise textures and objects within 37 minutes of data (i.e. 4 runs x ∼9-min per run). Specifically, in panel 6a and 6b we report the voxels that were significantly active above baseline for the t-contrast faces ∩ noise textures ∩ objects > 0 (t(702)=3.10; q<0.05 FDR-corrected), superimposed on a co-registered T1-weighted MP2RAGE^63^ anatomical reference. Fig. 6a shows statistically significant activation in the posterior cingulate cortex. The related timecourse for a single run (∼9 min) is shown as an insert in the middle panel of Fig. 6a; the timecourse was averaged across all voxels under the crosshair. Fig. 6b shows a statistically significant activation in different slices from those shown in Fig. 6a; these slices were selected to illustrate activation in the prefrontal cortex and the related timecourse for a single run, averaged across all voxels under the crosshair. By decreasing the t-threshold to t(702)=2.5 (p=0.012 uncorrected) we can detect activation bilaterally in the lateral geniculate nucleus (LGN) (Fig. 6c). The related single-run timecourse reflects the average of all significantly active voxels for the left LGN only.

**Fig. 6:**
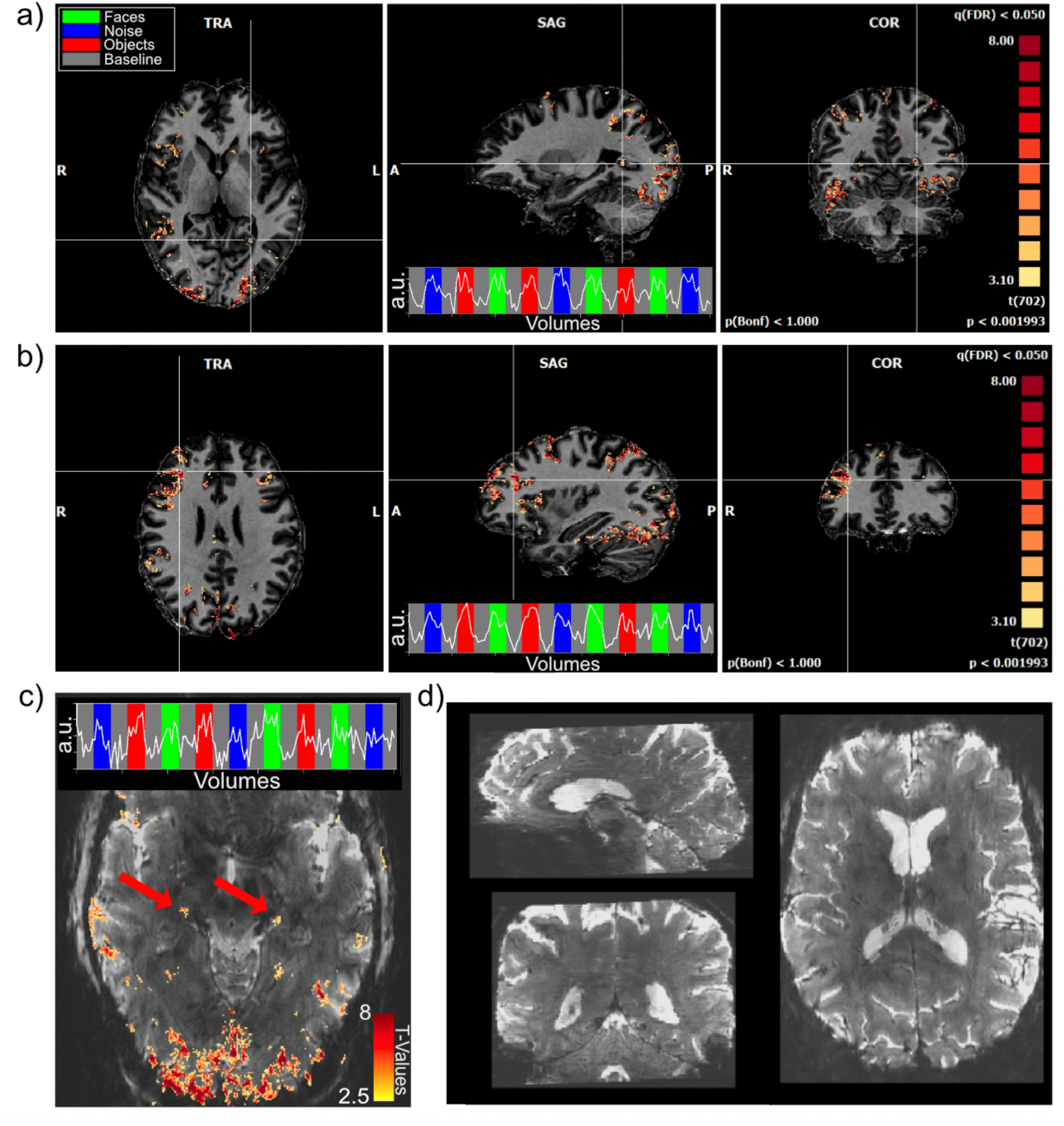
0.75 mm isotropic whole-brain fMRI activation (t-maps) elicited by a visual task: faces, noise textures and objects > 0, for approximately 37 minutes of data (without any denoising). **(a)** and **(b)** show activation maps in two different sets of three orthogonal slices (superimposed on co-registered MP2RAGE images) selected to show activation in the posterior cingulate cortex **(a)** and prefrontal cortex **(b).** The time courses averaged for all voxels in the cluster identified by the crosshairs are shown for a single ∼9-min run in the middle panel of both Fig. 6a and 6b. **(c)** activation in bilateral LGN (red arrows) after 2-voxel smoothing superimposed on a single EPI image and related time course in a single run. **(d)** Example sagittal, coronal and axial slices of the 0.75 mm isotropic whole-brain single EPI volume.

## 3. DISCUSSION

Understanding the function of distributed large-scale brain networks that underlie behavior ultimately requires information about the nature of the minimal architectural units that organize neural populations of similar properties. Such elementary cortical units exist in the mesoscopic scale, exemplified by complex but orderly organizations tangent to the cortical surface, such as the canonical orientation domains, and the cortical columns that span the multiple layers of the ∼1.5- to ∼4-mm-thick mammalian cortex. These meso-scale units are relevant also for untangling the mechanisms operative in pathological developments; for example, perturbations in layer-dependent computations that integrate feedforward, feedback and recursive circuital interactions have been proposed as models of dysfunction in neuropsychiatric disorders^64, 65^. However, despite notable advances demonstrating feasibility, the ability to perform fMRI with sufficient resolution and fCNR to robustly access these meso-scale computations remains suboptimal, leading, as previously mentioned, to the incorporation of ambitious resolution targets in the strategic plan of the US BRAIN Initiative that are far beyond existing capabilities. Achieving such resolution goals requires significantly higher image SNR, substantial elevation in the intrinsically weak fMRI signal changes evoked by neuronal activity, and the ability to counteract the increasing acquisition-time demands of high-resolution imaging. It also requires remedies to imperfections inherent in the fMRI method that degrade the neuronal specificity of the functional signals. We demonstrate that the new frontier of beyond-10T magnetic fields, in this case 10.5 Tesla-addresses these requirements and establishes a path forward for achieving major improvements in fMRI.

Imaging at such high magnetic fields, however, poses several challenges. These include heterogeneous transmit B1 (B1^+^) over the brain, larger B0 inhomogeneities that induce distortions in EPI acquisitions that remain the preferred acquisition method in fMRI, and signal fluctuations induced by physiological processes, mainly respiration. In this demonstration of high-resolution functional imaging at 10.5T, several strategies were used to overcome these challenges. All data processing incorporated dynamic off-resonance in k-space (DORK) processing that minimizes respiration induced noise^66^. 16-channel transmit capability was utilized to generate transmit B1 solutions that provided sufficient signal uniformity to obtain functional images over whole-brain or large sections of the brain (Fig. 5 and 6). The resulting signal uniformity can be seen in the GRE EPI images from which fMRI data are calculated and shown either by themselves (Fig. 5 and 6d) or together with functional maps derived from them (Fig. 3 and 4).

At high resolutions, where thermal noise dominates, the optimum GRE BOLD contrast occurs when TE (to the center of k-space) is ≅T2*. Achieving this optimal TE is another challenge faced in high resolution imaging in general, but especially at very high magnetic fields like 10.5T since the T2* decreases with increasing magnetic field strength. For high resolutions, segmentation of the EPI acquisitions to cover the in-plane k-space (i.e. typically designated as k_x_-k_y_) offers one solution to meet this TE requirement. However, such segmentation prolongs volume coverage and introduces increased susceptibility to physiological noise (e.g. ^67^). Therefore, we in general prefer avoiding this strategy. Using partial Fourier aggressively, if need be (e.g. 5/8), and high in-plane accelerations, we were able to achieve TE equal to or even less than the T2*. This was enabled by the inherently higher acceleration achievable at increasing magnetic fields as previously demonstrated for 10.5T for the RF coils employed here. ^20, 21^ In large FOV acquisitions, where reducing VAT also becomes imperative, we were able to achieve TEs that were less than the T2*. Although this deviates from the optimal condition, the BOLD contrast maximum with TE is relatively broad^5^; therefore, losses encountered vary slowly with the deviation from the ideal condition, and the loss can be compensated by the shorter TR and higher number of data points achievable within a given experimental time.

Finally, although B0 inhomogeneities do distort EPI images and this is more of a challenge with increasing magnetic fields, the problem is also alleviated by high in-plane accelerations available at such high magnetic fields with low g-factor noise. These distortions are a problem when accurate correspondence is sought for functional maps with anatomical features determined from other imaging approaches. However, this problem is avoided when the same GRE EPI images that form the basis of the fMRI time series also provide anatomical landmarks. We demonstrate this here in the primary visual cortex, thus avoiding image registration among different acquisition strategies. This is illustrated in Fig. 4 where a mid-cortical structure detected as hypointensity in MEGRE images (Fig. 1) was also detectable in the individual images of the GRE EPI fMRI time series and was used to align images for layer analysis directly in the fMRI space. There exists overwhelming evidence based on *in vivo* MR as well as *ex vivo* histology and MR studies that link this hypointensity in T2*-weighted MR images in the visual cortex to the Stria of Gennari ^68–74^, with iron accumulation thought to be the dominant source of this contrast ^70^. Besides histology, functional studies also confirm the assignment of this hypointense band to Stria of Gennari. The Stria of Gennari exists in visual area V1 (area 17) and ends at the V1-V2 borde*r*. The V1-V2 border can be identified using fMRI exploiting the retinotopic organization of these two regions and is also easily identified in the macaque brain based on the gyral/sulcal features. In the macaque fMRI experiments, the mid-layer hypointensity in T2*-weighted images detected in the V1 cortex was shown to terminate at the V1-V2 boundary^74^, a hallmark of the Stria of Gennari. The position of the Stria of Gennari corresponds to Brodmann layer 4B, or, in a different layer classification^75^ created to harmonize mammalian and rodent layers, to layer LIIIc, immediately adjacent to the input into V1 coming in layer 4C (Brodmann) or L4a, and L4b in reference ^75^.

When comparing fMRI properties at two different magnetic fields, as done in this paper, it is imperative to consider gains independent of hardware details *versus* those achieved with the current state of instrumentation. In fMRI, ultimately functional maps are determined by fCNR, which is given as Δ*S*/*N* where *N* is “noise” reflecting the fluctuations in the fMRI time series^27^ and Δ*S* is the signal change induced in the baseline signal *S* by the stimulus or task. This can be cast as *fCNR* = Δ*S*/*N* = (Δ*S*/*S*)(*tSNR*) where Δ*S*/*S* is fractional or percent signal change (PSC) coupled to neuronal activity, and *tSNR* is the temporal SNR in the fMRI time series. In this expression, Δ*S*/*S* is **independent** of hardware details such as the RF coils or gradients employed (see SUPPLEMENTARY Information, section 3). For GRE BOLD fMRI, 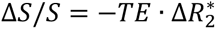, where 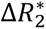 is the difference in 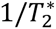 between the activated and control states in task/stimulus-based fMRI^5^ (SUPPLEMENTARY Information, section 3). 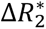 is a measure of the BOLD effect reflecting the mechanism underlying BOLD contrast. It depends strongly on vascular architecture, the neurovascular and metabolic responses to altered neuronal activity, and the magnetic field strength. We show that at equally optimal and the same TE for 10.5T and 7T, Δ*S*/*S* increases by a factor of ∼2 at 10.5T vs 7T for GRE BOLD fMRI, indicating that 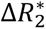 is a factor 2 higher at 10.5T.

Expressed as a power of the static magnetic field, a factor of 2 increase in Δ*S*/*S* from 7 to 10.5T reported here corresponds to a 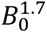 dependence. This is possible if the BOLD effect at 10.5T is dominated by the microvascular contributions. All modeling studies predict only a linear increase dependence on B_0_ for the large blood vessels (e.g.,^22, 23, 26^) while up to quadratic dependence has been predicted for the microvascular BOLD effect (e.g.,^22, 26^). Although more complex modeling studies of the total fMRI signal itself (as opposed to the BOLD effect alone) suggest a smaller increase for the microvascular contribution with increasing magnetic fields^23^, such integrated models rely on a large number of assumptions on physiological and metabolic changes associated with neurovascular coupling and vascular anatomy^23^ and may not reflect accurately the experimental conditions.

In contrast to Δ*S*/*S*, RF coils can play an important role in fCNR and tSNR because of their impact on image SNR. All coils fail to capture 100% of the theoretically maximal SNR (i.e. uiSNR). However, coils that capture the same fraction of uiSNR, i.e. have the same “SNR performance” at two different magnetic fields, can be employed for comparing functional maps and fCNR across magnetic fields. This was done for subjects 1 and 2 in data presented in Fig. 2 (see SUPPLEMENTARY Information, section 2). These data give an indication of the improvements expected at 10.5T relative to 7T, and demonstrate, all else being equal, significant gains in the ability to detect activation. These are PSC maps for voxels classified as active by a set threshold and show a larger number of activated voxels at 10.5T compared to 7T. On the other hand, S3 in Fig. 2 was obtained by using a different and new generation 64-channel receive 7T coil with higher SNR performance (SUPPLEMENTARY Information, Fig. S3). The difference between the 10.5T and 7T data in terms of number of voxels classified as activated is less dramatic, but nevertheless there. Functional maps at both magnetic fields can improve with better RF coils, especially in the peripheral regions where the uiSNR is intrinsically very high but RF coils in general capture a small fraction (∼20% or less) of the available uiSNR^76^ (also see Supplementary Figs. S2 and S3).

tSNR in the brain depends on image SNR but also can be strongly impacted by contributions originating from physiological/biological processes. It is a complex parameter that can be expressed as *tSNR* = *S*/(*N_T_* + *N_Br_* + *N_R_*), where the denominator represents temporal fluctuations in a time series with *N_T_* denoting instrumental thermal noise (ignoring instrumental instabilities, see SUPPLEMENTARY Information, section 1), *N_Br_* characterizing spontaneous oscillations associated with spatially correlated neural activity, the substrate for resting state fMRI, and *N_R_* representing B_0_ fluctuations by non-neuronal physiological processes such as respiration and heartbeat. *N_Br_* arises from the BOLD effect and hence increases with signal strength and the magnetic field. Similarly, respiration induced effects are proportional to B_0._^66, 77^ Even when *N_R_* is fully suppressed using various strategies (e.g.,^78–83^), it is possible that tSNR does not increase with magnetic field despite the substantial increase in image SNR. However, tSNR, as previously mentioned, is a highly complex parameter that depends on many factors including spatial location in the brain, different tissue types, acquisition details, and post-processing denoising strategies; such a full evaluation of tSNR is beyond the scope of this paper. Nevertheless, even in the limit tSNR does not change with increasing B_0_, 10.5T will still have a major advantage in task/stimulus-based fMRI because of the 2-fold gain in 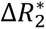 and Δ*S*/*S*.

However, in the limit *N_T_* ≳ (*N_Br_* + *N_R_*), which would be reached at sufficiently high spatiotemporal resolutions, *tSNR* gains at the higher magnetic field should also start contributing to fCNR. The 0.4 mm isotropic resolution GRE BOLD fMRI studies comparing 10.5T vs 7T presented in Figure 2 appear marginally to be in this regime (SUPPLEMENTARY Information, section 1). Higher spatiotemporal functional imaging can push the data well into the thermal noise dominated regime. In this limit, an upper bound on the *fCNR* gains possible at 10.5T can be calculated in the context of uiSNR, which is ∼2 to ∼2.5-fold higher over the brain at 10.5T compared to 7T ^19^. Together with the 2-fold increase measured in PSC, this yields a theoretically maximal gain of ∼4 to ∼5-fold higher fCNR in GRE BOLD-fMRI over the brain at 10.5T vs 7T. Experimentally, SNR at 10.5T appears sufficient to reach resolutions that fall in this regime, especially with additional developments in RF coil performance; however, such higher resolutions will face another challenge, namely image acceleration. Solutions to this challenge will be acceleration optimized RF coils, better image reconstruction, and high-performance gradients.

Higher spatial resolutions require longer EPI echo train length (*ETL*), and signal decay during the longer *ETL* leads to blurring, especially at higher magnetic fields where T2* gets shorter. As such, pushing to higher nominal resolutions with EPI-based fMRI can in fact lead to a decrease in effective resolution^84^. This problem can be countered using high-performance gradients^84^ and/or a high degree of in-plane parallel imaging; in both cases, there is an SNR loss, which is the same for the same reduction factor in *ETL* (see SUPPLEMENTARY Information, section 4). This loss comes from the higher bandwidth needed with high-performance gradients for reducing *ETL* and the fewer k-space data points collected with parallel imaging. The latter is limited by the spatially non-uniform noise amplification given by the g-factor^85^. However, ultrahigh fields (e.g.^21, 86, 87^) and high channel count arrays (e.g.^21, 88, 89^), as employed in this study, are particularly effective in suppressing the g-factor and promoting high accelerations. With this approach, we demonstrate that at 10.5T, even though there exists blurring, we are indeed gaining *effective* resolution when the *nominal* resolution is increased to 0.35 mm isotropic (Fig. 3 and 4). In contrast, at 7 Tesla, blurring precluded gains in resolution beyond ∼0.6 mm when using in-plane parallel imaging acceleration of 3 and standard gradient performance equivalent to that employed in the 10.5T at present.^84^

Depending on the region and the paradigm used, laminar profiles of functional signals should in principle display a peak in the middle layers for feedforward-dominated tasks (e.g.^90–92^) or two peaks associated with deeper and superficial layers for feedback-dominated tasks^93^. Instead, progressively increasing PSC towards the pial surface is a renowned laminar feature of GRE BOLD fMRI (e.g.^90, 91, 93, 94^), as also illustrated by the 0.8mm 7T data in Fig. 4b. This is caused by the strong BOLD responses originating from the large draining veins at the pial surface, which extend into the cortex and decrease with increasing distance from the pial surface. Nevertheless, due to its superior sensitivity, temporal resolution, and coverage, GRE BOLD remains the most popular acquisition protocol for fMRI, even for meso-scale studies, where the limited specificity is a real problem. Differential mapping can ameliorate, although not necessarily eliminate this problem, but requires suitable paradigms. Acquisition techniques that encode spin-echo BOLD contrast such as SE EPI (e.g.^90, 92, 95–97^) or SSFP ^98–100^ can suppress the large vessel contributions but do so at the expense of a decrease in BOLD effect and fCNR due to diffusion averaging (e.g. ^101, 102^, also see SUPPLEMENTARY Information, Fig. S5). VASO ^28, 103^, a technique that reports on cerebral blood volume changes and is largely immune to the large vessel confound, also suffers from decreased SNR and fCNR^28, 103, 104^. Here we show that ultrahigh resolutions (0.35mm nominal) achieved at 10.5T for *single condition* GRE BOLD responses in fact suppress the draining-vein confound, as we predicted previously^56, 105^, yielding accurate cortical layer profiles. The evidence for this comes from several observations.

First, we observe a decrease in the slope of the layer profiles of the 0.35mm 10.5T fMRI data vs. the 7T 0.8mm images (SUPPLEMENTARY Information, Fig. S8). Second, we can clearly distinguish activation across adjacent banks of a sulcus for the 0.35mm 10.5T data but not for the 0.8mm 7T images (Fig. 3c and Fig. 4c). And finally, at 0.35mm, we can detect *features* across cortical layers that are not visible with standard 0.8 mm resolution GE BOLD images. Specifically, we observe the presence of a mid-cortical maximum in PSC, appearing as a “bump” in the laminar profile of the 10.5T 0.35mm data, even in a ∼5-min run. This maximum corresponds to the location of the Stria of Gennari and therefore to the middle layers. As demonstrated by invasive animal studies, V1 receives direct feedforward input from the LGN into middle layers. As such, the larger activation in middle layers is precisely what is expected from pure visual feedforward stimulation as implemented here.

There are two reasons for this improved specificity. One is a magnetic field effect, namely the supralinearly increasing microvascular contribution at the higher magnetic fields. The second is a resolution effect, the existence of which was discussed in our previous publications^56, 105^ and is more extensively presented here in the SUPPLEMENTARY Information Section 6. The extravascular BOLD (EV-BOLD) phenomenon dominates BOLD-based fMRI maps at ultrahigh magnetic fields (e.g.^23 106^); it generates fMRI signals through the intra-voxel dephasing of spins due to magnetic field inhomogeneities induced outside deoxyhemoglobin containing blood vessels. In the limit voxel dimensions become small compared to the distances that characterize these magnetic field gradients associated with EV-BOLD, magnetic fields *inside* the voxels become more uniform, suppressing the BOLD response. This is particularly true with increasing distance from the blood vessel as the magnetic field gradient becomes shallower. A prediction of this effect is that the BOLD effect associated with pial veins shrinks in spatial extent, persisting only in close proximity to the blood vessel boundaries where the magnetic field gradient is steepest (see SUPPLEMENTARY Information Section 6). Thus, as resolutions increase, draining-vein effects get concentrated at the pial surface, and penetrate less into the cortex. This is indeed what is observed in the 0.35mm data where the PSC rapidly rises just beyond the outer cortical boundary in the space where the pial veins reside (Fig. 4b). In principle, even at 7 Tesla, where BOLD effect originating from microvasculature exists, such high resolutions should produce the same effect seen in Fig. 4a, although this is unlikely in just a 5 min acquisition since fCNR will be smaller.

This resolution effect can also be conceptualized using simple GRE images of a brain. It is well recognized that intravoxel dephasing reduces with smaller voxel sizes and this effect is reproduced in Supplementary Fig. S6.2 where signal intensity dropouts are visualized near the air-filled cavities at low but not high resolution.

It would be ideal to use such high-resolution data for defining brain-wide networks with layer specificity, using Human-Connectome-Project (HCP) style whole-brain acquisitions. However, even with the high accelerations available from 10.5T and high channel count arrays, nominal resolutions such as 0.35 mm lead to undesirably long volume acquisition times. Nevertheless, compromises are possible that provide significant improvement over the 1.6 mm resolution 7T HCP resting-state and task-based fMRI acquisitions^61, 107^. Here, we demonstrate the feasibility of next generation HCP style high-resolution, whole-brain fMRI, one that can achieve sub-millimeter (0.75 mm) resolutions with TRs of 3s, while maintaining sufficient functional sensitivity. We used 12-fold 2D acceleration in a GRE SMS/MB EPI acquisition, exploiting combined through-plane (i.e. multiband, MB^60^) and in-plane acceleration and partial Fourier to decrease TE and TR, thus reducing the image acquisition times to practical levels. Single-subject functional images obtained in ∼37 min with this protocol display robustly detectable activity evoked by faces, noise textures and objects, extensively through the brain, ranging from LGN, visual cortex, temporal lobes, parietal lobes, prefrontal cortex, etc. (Fig. 6). Functional mapping at these spatiotemporal resolutions and coverage in single subjects represents an unprecedented opportunity for human neuroscience.

## 4. CONCLUSION

In conclusion, we present initial results from 10.5T GRE BOLD-fMRI demonstrating major increases in fCNR, achievable resolution, and spatial precision of functional mapping, exploiting the major gains in intrinsic SNR and image acceleration available for brain studies at 10.5T. We also demonstrate that high spatial resolutions and high magnetic fields suppress the draining-vein confound of the GRE BOLD-fMRI, improving its specificity relative to neuronal activity. The results presented usher in the gains available in the new era of beyond 10 Tesla human functional imaging, particularly for meso-scale fMRI studies. Further improvements in effective resolutions and temporal sampling (i.e. acceleration) are possible when such extremely high fields will be combined with the use of higher performance gradients, better performing RF coils, and further developments in image reconstruction and denoising strategies, such as those based on deep learning (e.g.^108, 109^).

## 5. METHODS

### 5.1. Experimental paradigms

#### 5.1.1. Block design retinotopically organized visual stimulation with flickering gratings

The experimental procedure consisted of a 24-second on/off block design for the 0.4 mm isotropic 7T/10.5T acquisitions and for the 10.5T 0.35 mm isotropic voxel acquisitions. For the 7T 0.8 mm isotropic resolution dataset the durations were 12s on and 12s off for S3. The difference in block length was implemented to account for the difference in volume acquisition time between the 0.8 mm and the other acquisitions (see later). For the remaining subjects we acquired the 7T 0.8 mm images with identical timings as the 10.5T 0.35 mm images (i.e. 24 seconds on/off). The stimuli consisted of a center (i.e. target) and a surround square checkerboard counterphase flickering (at 6 Hz) gratings subtending approximately 5.6 degrees of visual angle. Stimuli were centered on a background of average luminance (25.4 cd/m^2^, 23.5-30.1). Using a Mac Pro computer, stimuli at 7T were presented on a Cambridge Research Systems BOLDscreen 32 LCD monitor positioned at the head end of the scanner (resolution 1920, 1080 at 120 Hz; viewing distance ∼89.5 cm). At 10.5 T, stimuli were presented using a VPIXX projector (resolution of 1920 × 1080 pixels at 60 Hz; viewing distance ∼89.5 cm). Stimulus presentation was controlled using Psychophysics Toolbox (3.0.15) codes. Participants viewed the images through a mirror placed in the head coil.

Each run began and ended with a red fixation dot centered on a gray background and consisted of 3 trials for each of the 2 conditions. This resulted in eight ∼2.5-minute runs for the 12 seconds on/off acquisition (i.e. S3’s 0.8 mm 7T data) and just over 5-minute runs for the 24 seconds on/off acquisitions (6 runs). Participants were instructed to minimize movement and keep fixation locked on the center dot throughout the experimental runs.

#### 5.1.2. Block design visual stimulation with faces, noise textures and objects

The experimental procedure implemented for the whole-brain isotropic 0.75 mm acquisition consisted of a standard 24-second on/off block design, where with “on” we refer to blocks with visual stimulation, while for “off” we refer to fixation periods (i.e. no visual stimulation). The stimuli consisted of a set of face images, noise textures, and objects (i.e., on-blocks), subtending approximately 5° of visual angle. We equated the amplitude spectrum across all images. The stimuli were centered on a background of average luminance (25.4 cd/m^2^, 23.5-30.1) and were presented using a Mac Pro computer and a VPIXX projector (see above). Stimulus presentation was controlled using Psychophysics Toolbox (3.0.15). Participants viewed the images through a mirror placed in the head coil.

Each run began and ended with a 24-second fixation cross centered on a gray background (i.e., off-block) and consisted of 3 trials for each of the 3 conditions (∼7 and a half minutes duration). We collected 4 experimental runs. Participants were instructed to minimize movement and keep fixation locked on the center fixation dot throughout the experimental runs. This experimental paradigm was adapted from ^110^.

#### 5.1.3. Auditory stimulation

For both acquisitions, we adapted (using cascading sequences) the classical local-global paradigm that has been used previously to study the effects of local and global expectations on ‘error-related’ responses ^111^. Each trial consisted of a sequence of 4 sounds (100 ms per sound with a 100 ms gap in between) that was presented over a total of 700 ms, followed by a 500 ms gap before the next trial started. For each sequence, the first three sounds were presented in an ascending or descending order in acoustic frequency with the fourth tone either following this ascending or descending pattern (locally standard trials) or violating this pattern (locally deviant trials). In distinct experimental blocks of 3 minutes each, global regularity was defined according to the presentation rate of each stimulus. Globally standard trials were presented 90% of the time, whereas globally rare stimuli were pseudo-randomly presented 10% of the time. Four types of trials were created by manipulating local and global regularities: local and global standard, local and global deviant, local standard but global deviant, and local deviant but global standard.

To account for the presentation rate of each frequency, in each block we presented two types of globally standard trials (e.g. frequencies [1 2 3 4] and [2 3 4 5]) that were presented pseudo-randomly. In between globally deviant trials (e.g. frequencies [2 3 4 1]), we always presented either 8, 9 or 10 globally standard trials. Each block started with 30 habituation trials (not counted towards the 90%-10% regularity rule) to establish the global regularity before presenting the first globally deviant trial. Each run consisted of two blocks, one in which the globally standard trials were also locally standard (e.g. [1 2 3 4] and [2 3 4 5]) and alternatively one block in which globally standard trials were locally deviant (e.g. [2 3 4 1] and [3 4 5 2]) and the globally deviant trials were locally standard (e.g. [2 3 4 5]). The order of blocks, and the ascending and descending cascades were counterbalanced. Each run lasted approximately 8 minutes and we recorded a total of 8 experimental runs.

### 5.2 MR Imaging Acquisition and Processing

The 0.8 mm^3^ 7T functional MRI data were collected with a 7T Siemens Magnetom System with a single transmit and 32-channel receive NOVA head coil, while the 0.4 mm^3^ were collected with a 7T Terra System with an in-house-built single-channel transmit and 64-channel receive head coil ^19, 20, 112^ and a new 64-channel receive, 16-channel transmit array built by MR CoilTech, Ltd (UK). All 10.5T MRI data were collected with a 10.5T Siemens Magnetom System with an in-house built 16 transmit and 80 receive -channel head coil ^19, 20, 112^. The in-house developed 10.5T and 7T array coils have been extensively characterized in terms of performance ^19, 20, 112^.

All 10.5T studies and the 7T study with the 16-channel MR CoilTech transmit coil utilized pTx with amplitude and phase shimming, i.e. single-spoke pulses; for whole-brain studies, a solution optimizing 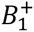 for the whole-brain was sought. For studies of the visual cortex using a limited coverage, the optimization was performed over a restricted FOV which included but was larger than the volume that was covered by the imaging slab. In both cases, subsequent to the pTx pulse generation, the flip angle was adjusted over the volume/slab of interest.

Signal intensity in a GRE-EPI-based fMRI experiment is affected by 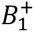 but also T1 and TR since full relaxation is generally avoided; this interaction together with reduction in the spatial variation in 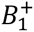 tends to reduce spatial variations in the resultant signal intensity in the image. This was the case for the GRE-EPI images used for fMRI data both for whole brain (Fig. 5 and 6) and restricted field-of view (Fig. 3). In addition, fMRI in general is not so sensitive to 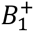 provided the voxel has enough SNR because fMRI looks at changes in the signal intensity. In particular, as discussed extensively in section 3 of the SUPPLEMENTARY Information, PSC, a major focus of this paper, is independent of 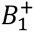 and 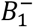.

#### 5.2.1. 10.5T acquisition protocols

Below we list the acquisition parameters for all protocols:

##### 0.65 mm isotropic nominal resolution fMRI

*3D GRE EPI;* TR=81 ms; TE=16.6 ms; in-plane acceleration factor along the phase-encode direction (iPAT)=4; PF=5/8; 52 Slices; VAT=4536 ms; FA=14deg; Matrix size=276×272×56; bandwidth=954 Hz/Px; slice oversampling 0%; ETL=43; echo spacing=1.19 ms.

*2D GRE EPI*; TR=2320ms; TE=20.6 ms; iPAT=4; Multi Band=2; PF=5/8; 60 Slices; FA=67deg; Matrix size=291×296×60; bandwidth=1008 Hz/Px; ETL=47; echo spacing=1.17 ms.

##### 0.4 mm isotropic nominal resolution fMRI

3D GRE EPI; TR=111 ms; TE=24.2 ms; iPAT=3; 5/8 PF; 40 Slices; VAT=4884 ms; FA=15deg; Matrix size=364×220×40; bandwidth=654 Hz/Px; slice oversampling 10%; ETL=46; echo spacing=1.68 ms.

##### 0.35 mm isotropic nominal resolution fMRI

3D GRE EPI; TR=113 ms; TE=24 ms; iPAT=4; 5/8 PF; 40 Slices; VAT=4972 ms; FA=15deg; Matrix size=364×260×40; bandwidth=598 Hz/Px; slice oversampling 10 %; ETL=41; echo spacing=1.91 ms.

##### 0.75 mm isotropic nominal resolution fMRI

2D SMS/MB GRE EPI; TR=3210 ms; TE=17.80 ms; iPAT=4; Multi Band=3; 5/8 PF; 144 Slices; bandwidth=1068 Hz/Px; Matrix size=260×272×144; FA=75deg; ETL=43; echo spacing=1.06ms.

##### MP2RAGE Anatomical imaging

0.7 mm isotropic voxels; TR=5000 ms; TE=1.97 ms; Acquisition time: 07:30 minutes; FOV=253*270mm; iPAT=4; Inversion 1=840 ms; Inversion 2=2370 ms; Partial Fourier 6/8; FA 1=6deg; FA 2=6deg

*<u>Multi Echo</u> Gradient-Recalled Echo (MEGRE) <u>acquisition with 0.37 mm isotropic resolution</u>*: to estimate T2* values, whole brain data on two subjects were obtained with a 3D MEGRE acquisition with 7 echoes: TEs: 4.07, 7.95, 11.83, 15.71, 19.59, 23.47, 27.35 ms; TR=33 ms; Acquisition time: 9:44 minutes. In the following acquisitions geared towards identifying the Stria of Gennari rather than estimating T2* values, we only acquired 6 echoes (TEs; 4.1, 7.9, 12, 16, 20, 23ms; TR=29ms) to shorten the acquisition time (8:33 minutes). The remaining acquisition parameters were kept constant: FA=10deg; 7/8ths partial Fourier; 2D acceleration (in-plane and through-plane) GRAPPA: 3×3; Matrix (*k_x_* × *k_y_* × *k_z_*)=576×576×320, with *k_z_* designating the through-plane “slice” direction.

##### MEGRE acquisitions 0.8 mm isotropic resolution

Whole-brain data on three subjects were obtained with 3D MEGRE using parameters: TEs: 4, 8, 12, 16, 20, 24, 28 ms; TR=33 ms; FA=10deg; 7/8ths partial Fourier; GRAPPA: 3×2; Matrix(*k_x_* × *k_y_* × *k_z_*)=264×264×240 with *k_z_* designating the through-plane “slice” direction. Acquisition time: 5:34 minutes.

#### 5.2.2. 7T acquisition protocols

##### 0.4mm isotropic nominal resolution fMR

3D GRE EPI; TR=113 ms; TE=24.4 ms; iPAT=3; 5/8 PF; 40 Slices; VAT=4972 ms; FA=16deg; Matrix size=368×220×40; bandwidth=654 Hz/Px; slice oversampling 10%, ETL=46; echo spacing=1.67 ms.

##### 0.8 mm isotropic nominal resolution fMRI

2D GE EPI; TR=1350 ms; TE=26.4 ms; Multi Band=2, iPAT=3; 6/8ths Partial Fourier; 42 Slices; FA=70deg; Matrix size=210×198×42; bandwidth=1190 Hz/Px; ETL=50; echo spacing=1 ms.

##### MEGRE acquisitions 0.35 mm isotropic resolution

Six echoes, partial field of view (axial slab, roughly aligned to AC-PC line) multi-echo 3D GRE images were obtained using a 32 Channel NOVA head coil. TEs 4.37, 8.74, 13.11, 17.48, 21.85, 26.22ms; TR=32ms; FA=12deg; no partial Fourier; GRAPPA 2×2. Matrix(*k_x_* × *k_y_* × *k_z_*)= 576×576×160 with *k_z_* designating the through-plane “slice” direction. Acquisition time: 10:18 minutes.

### 5.3 MEGRE data processing for calculation of T2* values

#### Processing

Due to the bipolar readout, visible distortion was present in odd versus even echoes at 10.5T. To correct for this, we calculated nonlinear alignments between the odd and even echoes (correcting to the intermediate space) using the SyN approach in the antsRegistration package. To align across runs, both rigid and nonlinear alignment was calculated, using the same package. A final alignment step to the anatomical space from which regions of interest were defined (described below) was also calculated using the same tool. All transforms were combined and applied in a single step to prevent blurring from multiple interpolation steps. These steps follow current best practices for the analysis of such high-resolution data^57^.

Following the transformation, a grand mean was calculated for each echo across the 2 or 3 separate runs.

For the 7T data, distortions were minimal in our regions of interest, and therefore nonlinear methods were not used. Instead, the mean of each run was rigidly aligned to the first run and then to the anatomical space. Following the transformations, a grand mean was calculated for each echo across the 6 separate runs. Finally, both the 7T and 10.5T data were aligned into the same space for comparison.

#### T2* Fits

T2* fits were estimated using the function t2smap from the python package tedana, v23.0.3 (https://github.com/ME-ICA/tedana) using the nonlinear curve fitting approach, which is slower but expected to be more accurate than the typical log-linear approximation. The values in Fig. 1c are derived from the weighted average across echoes, optimizing the contrast for each voxel based on the T2* fits also shown in Fig. 1c, restricted to V1, out to an eccentricity of 90 degrees (from Benson 2014 atlas). That is, based on the fits, the voxels were combined to weight the resulting image towards TE = estimated T2*. Fig. 1c left shows the voxel values after this process, while the right shows the corresponding voxel estimated T2* values. The reason for using the raw average versus weighted average in Fig. 1a, was to show the entire brain (the weighted average requires brain masking).

#### Regions of Interest

A mask of the gray matter (GM) ribbon was produced at high resolution (0.37mm) using the surfaces (smoothwm, pial) derived from FreeSurfer version 7.2.0. Labels for white matter (WM) and insula areas were derived from the automatic labeling performed by Freesurfer, after upsampling to the 0.37mm space. The label for V1 was produced using the Benson Atlas automated label approach^57^. For GM/WM comparisons a final inclusion mask was manually drawn to include only areas with sufficient signal in the 0.37mm 10.5T data. To produce cortical depths, the high-resolution GM ribbon, along with WM and CSF labels were passed through LN2_LAYERS - a part of the LayNii toolbox (version 2.7), using equivolume depth calculations. The metric file, which contains a cortical depth estimate for each voxel within the GM, was then used to extract depth-dependent metrics from the weighted average and T2* estimates.

### 5.4 Functional data Preprocessing

All functional data preprocessing was performed using BrainVoyager (version 21.4), AFNI (version 22.3.07) and ANTs (version 2.3.5). Preprocessing was kept at a minimum and consistent across reconstructions. Specifically, we performed slice scan timing corrections for the 2D data only (temporal sinc interpolation), 3D rigid body motion correction (spatial sinc interpolation) where all volumes for all runs were motion corrected relative to the first volume of the reference run, and low-frequency drift removal (i.e. temporal high pass filtering) was performed using a GLM approach with a design matrix continuing up to the 3^rd^ order discrete cosine transform basis set. No spatial or temporal smoothing was applied. Anatomical images (MP2RAGE and MEGRE) were aligned to a reference run of the functional data (mean across the timeseries) using AFNI and ANTs. For the ultrahigh resolution (0.35 mm and 0.4 mm) data, this involved both linear and non-linear transforms, manual adjustments and iterative optimizations. For the 0.65 mm and 0.75 mm data, linear transforms in combination with top-up distortion correction (using a reverse phase encode acquisition^113^) were found sufficient for accurate registration.

To quantify the extent of stimulus-evoked activation, we performed standard GLM estimation (with ordinary least squares minimization) using BrainVoyager or AFNI. Paradigm regressors of the design matrices were generated by convolution of a canonical hemodynamic response function with a “boxcar” function (representing onset and offset of the stimuli). Prior to GLM estimation, each voxel time series was scaled by its mean across all volumes of the run to a baseline value of 100, yielding regression coefficients for the task regressors expressed in units of percent signal change (%). Subsequent analyses (i.e. ROI-based and statistical testing) were performed in MATLAB using a set of tools developed in-house.

### 5.5 ROI definition and cortical depth segmentation

#### 0.4 mm isotropic voxel images

We further performed region of interest (ROI) analyses to compare evoked percent signal changes (PSC) across all voxels within the ROI. To avoid biasing the ROI definition toward either field strength, we defined our cortical region anatomically, guided by functional contrast maps. Specifically, the ROI was derived as follows: After co-registering both the 10.5T and the 7T functional data to the same MP2RAGE image, to identify the retinotopic representation of both stimuli in V1 (localized using Benson’s probabilistic atlas, see above) we performed the contrast target & surround > 0 of run 1 independently for each field strength (which was excluded from subsequent analyses). We used both t-as well as PSC maps at different thresholds across field strengths to maximize overlap between evoked activation. Guided by the contrast maps, we then relied on the Benson probabilistic atlas to identify V1 and proceeded to draw an anatomically informed ROI based on V1 delineations on the co-registered T1 image.

GLM t-values can be approximately conceptualized as beta estimates divided by GLM standard error according to this equation:

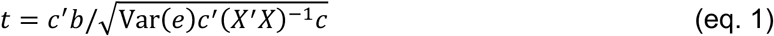

where b represents the beta weights, c is a vector of 1, −1 and 0 indicating the conditions to be contrasted, e is the GLM residuals and X the design matrix.

#### 0.8 mm and 0.35 mm isotropic voxel images

The ROI used for laminar analyses was defined in a portion of V1 corresponding to the retinotopic representation of the target condition. V1 was identified based on a clear manifestation of the Stria of Gennari in the 0.35 mm resolution mean EPI and MEGRE images. The target representation was selected as a patch of cortex demonstrating robust target-preference based on the target > surround contrast maps (p<.05 uncorrected). Note that the first two runs were used for ROI definition and thus discarded in subsequent laminar analyses. Single voxel time courses displayed in Fig. 3 were randomly selected from among significantly active (p<.05) voxels within the functionally defined retinotopic representation of the target stimuli in V1.

Voxel-wise cortical depth estimates were obtained using LAYNII’s LN2_LAYERS^114^, based on manually-drawn WM and CSF boundaries. These were drawn around the ROI on an upsampled (sinc interpolation) 0.2 mm isotropic grid. Laminar profiles were then computed by averaging the single-trial PSC estimates (target > fixation) of ROI voxels within each of 13 equivolume depth bins. It should be noted that partial voluming between the WM and GM as well as between GM and CSF, together with the interpolation method and the number of bins used, can impact edge regions and consequently laminar profiles. However, these effects are less pronounced at the current resolution, which approaches the upsampled resolution (0.35 mm vs 0.20 mm), providing a wider dynamic range for segmentation and minimizing partial voluming.

For S3, the number of trials used in activation maps and laminar profiles was 12 and 24 for the 0.35 mm and 0.8 mm datasets, respectively, to thereby match the total functional scan time.

Gradient-echo BOLD is plagued by contamination of unwanted venous BOLD, resulting in increased PSC in proximity of the pial surface (where large draining vessels are concentrated), giving rise to the renowned GE BOLD laminar ramping profile. To estimate the impact of draining vessels across resolutions, we quantified the slope of the amplitude ramping profiles across depths as follows: we performed a linear fit to the laminar profiles for each trial and resolution and estimated the slopes of the fitted lines using MATLAB’s *regress* function. Two-sample t-tests (across single-trial slope estimates) were used to evaluate for within-subject differences in slopes between the two resolutions. Note that for S3 only the first 12 trials in the 0.8 mm dataset were used in this t-test to match sample size across resolutions.

To assess statistical significance of the functional activation peak at the depth of the Stria in laminar profiles (Fig. 4b), we applied the following procedure independently per subject. For every trial, we fit a model consisting of a second-order polynomial (capturing the characteristic ramping profile of GE-BOLD), plus a Gaussian term (capturing the localized activation peak). Fitting was performed using MATLAB’s *lsqcurvefit* function (MathWorks Inc.). The model comprised three polynomial parameters (intercept p_1_, slope p_2_, quadratic term p_3_) and three Gaussian parameters (amplitude p_4_, mean p_5_ (depth-position), standard deviation p_6_ (width)). To prevent physiologically implausible solutions arising from overfitting—such as a monotonic decreasing polynomial compensated by an inflated Gaussian peak—we employed a leave-one-out fitting strategy. Specifically, for each trial, the polynomial component was first fitted to the mean profile of all remaining trials (polynomial-only model). The resulting slope (*p₂*) and quadratic term (*p₃*) were then fixed in the subsequent full model fit (polynomial + Gaussian; eq. 2) for the held-out trial. This procedure reduced model flexibility and was physiologically motivated by the assumption that the overall ramping shape of the GE-BOLD profile is largely constant across trials.

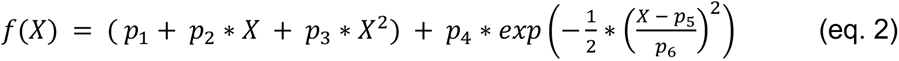

The intercept, slope and quadratic term were unconstrained. The Gaussian mean was constrained to the range of depth bins (1-13), and its standard deviation was constrained to the range corresponding to FWHM=2-6 bins to avoid modeling of physiologically implausible peak widths. The Gaussian amplitude, which constituted the parameter of interest, was unconstrained; the activation peak was considered statistically significant when this parameter was greater than zero, evaluated using a one-sample, one-sided t-test across trials. This directional choice was motivated by extensive prior literature establishing a strong *a priori* expectation of a positive activation peak at middle cortical depths^90, 92, 115–117^; deviations in the opposite direction would therefore constitute evidence against, rather than support for, the ability of the present setup to resolve feedforward layer-IV activity.

### 5.6 Quantifying BOLD images smoothness

Global smoothness measures were obtained to compare the estimated differences in functional blurring between different resolutions (0.35 mm versus 0.8 mm). Smoothness estimates were obtained from the GLM residuals, using the AFNI function 3dFWHMx^118^ with the ‘-ACF’ option. This method estimates the spatial autocorrelation from the data using a Gaussian plus mono-exponential model, which accounts for possible long-tail spatial autocorrelations. The estimated FWHM in mm from this fitted autocorrelation function is used as an estimate of the smoothness of the data. This estimate was derived from individual runs. Across-run paired-sample t-tests for the reconstruction comparison, and across-run two-sample t-tests for the resolution comparison were carried out between estimated FWHM parameters to infer statistical significance.

### 5.7 Tailored off-line reconstructions

The 3D-EPI was obtained with monotonic sampling of the slice-phase encoding direction and preceding each EPI readout a 3-line navigator was acquired without any phase-encoding. Each temporal multi-channel data sample was decorrelated based on the correlation estimate from a noise-only acquisition. Each EPI readout train was corrected for timing precision using a linear phase obtained from the 3-line navigator, with the same correction applied for all channels. Phase-encoding undersampling was corrected using a two-step GRAPPA implementation, where firstly the missing lines were calculated, and then the acquired data were re-estimated from the neighboring measurements, analogous to a 1-step SPIRIT. Ramp-sampling correction of each readout line was applied using a spline interpolation to the Nyquist sampling. A 1D FFT was applied in the slice-encoding direction. POCS partial Fourier reconstruction ^119^ was applied for each channel and each slice, and the reconstructed images were combined using a SENSE-1 algorithm with ESPIRIT estimated sensitivity profiles. NORDIC denoising was applied to the complex-valued time-series as described in ^120^, where the g-factor and noise were estimated using an implementation of the MPPCA algorithm.

### 5.8 Participants

We acquired 17 functional and 7 structural data sets on 7 healthy right-handed subjects (age range: 28-50), with different stimulation paradigms, acquisition parameters and field strengths (see *above*). All subjects had normal or corrected vision and provided written informed consent. The study complied with all relevant ethical regulations for work with human participants. The local IRB at the University of Minnesota approved the experiments.

## 5.9 Acknowledgments

This work was supported by NIH grants NIH 1R01NS136490-01 (X.W and L.V.) NIH U01EB025144 (K.U.); NIH P41 EB027061 (K.U.), NIH R01 EB038654 (A.S.) and NIH UM1NS132207 (K.U. and E.Y.)

## SUPPLEMENTARY Information

### 6.1. tSNR, the temporal signal fluctuations in the fMRI time series at 10.5T and 7T

tSNR as measured in the brain is a complex parameter that depends on many factors including spatial location, acquisition details, different tissue types, and post processing to suppress thermal (e.g. ^1^) and/or non-white, structured noise that exists in human brain images (e.g. ^2–8^). It also can vary for different subjects. As such, doing justice to magnetic field dependence of tSNR requires a detailed and comprehensive study, which is beyond the scope of this manuscript, where we focused on the B_0_ dependence of the BOLD effect quantified as percent signal change (Δ*S*/*S*). Nevertheless, we provide some tSNR data relevant to the field comparison given in this manuscript.

**Figure S1.1:**
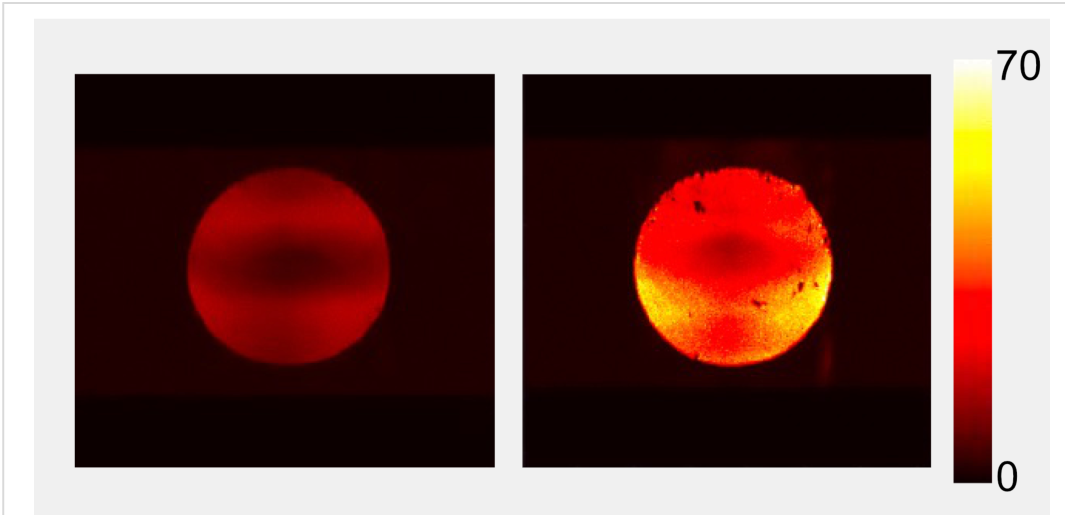
tSNR maps of a slice obtained in a phantom mimicking the average electrical properties of the and the of the human head and neck shape

tSNR can be degraded due to instrumental instabilities which include RF coil performance. This contribution cannot be evaluated in the human brain. Therefore, tSNR data for 0.4 mm isotropic resolution GRE EPI were acquired in a phantom using the experimental parameters employed for Figure 2 in the paper where 7T and 10.5T fMRI data are compared with respect to percent signal change associated with the BOLD response.

Figure S1.1 shows a tSNR map in a single slice obtained in a phantom (mimicking average electrical properties and the dimensions of the human head and neck; phantom details can be found in ^9–11^). Because of spatial variations in SNR over the sample, characteristic of high magnetic fields and array coils, we quote tSNR averaged over the slice (mean±std): tSNR was ∼2.6 times larger at 10.5T than 7T (31.23± 0.49 (std) for 10.5T *versus* 12.11±0.48 (std) for 7T). These numbers were calculated over ten consecutive ∼4 min runs and the standard deviation given is for the variation of the slice averaged tSNR for the ten separate runs. Correcting for the T2* decay during TE (24.2 and 24.4 ms for 10.5 and 7T, respectively) because the T2* of the phantom is field dependent (135 ms at 10.5T and 200 ms at 7T) yields an “intrinsic” 10.5T/7T tSNR ratio of 2.75. A flip angle correction was not applied; the flip angle was set to maximize the signal from the slab for the given TR for both magnetic fields.

**Figure S1.2.**
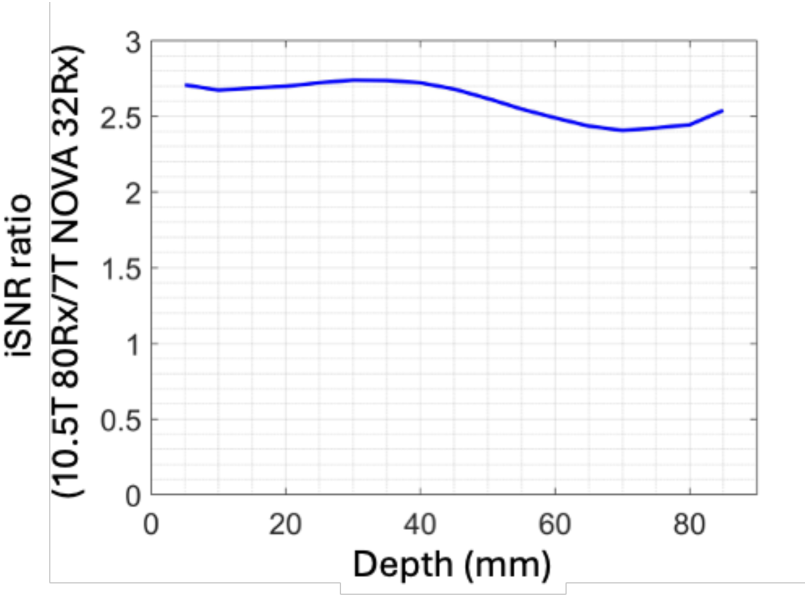
The ratio of intrinsic SNR (iSNR) measured at 10.5T with a CMRR built array with 80 receive channels *versus* 7 Tesla using the industry standard NOVA with 32 receive channels. iSNR reflects

In the phantom, the tSNR measured at two different magnetic fields (i.e. 10.5T and 7T in this case) should reflect SNR gains when temporal fluctuations in signal intensity arise predominantly from the thermal noise of the electronics. The magnetic field (10.5T vs 7T) and RF coil dependent SNR in this phantom have been documented for the coils employed here^11^ (industry standard NOVA 32 with 32 receive channels for 7T and CMRR built array with 80 receive channels for 10.5T). The ratio of intrinsic SNR (iSNR) at 10.5T vs. 7T as measured by these coils is shown in Figure S1.2 as a function of depth, measured as an average over concentric shells, outer most shell conforming to the contour of the phantom (see ^9–11^ for details). The iSNR ratio, 10.5T relative to 7T, was approximately 2.6 averaged over the different depths (Figure S1.2). This reflects field and coil dependent SNR gains; the former scales approximately quadratically with B_0_ over central regions, and the 10.5T coil performs better than the 7T coil in capturing the theoretically maximal SNR available at the respective magnetic fields^11^.

The iSNR ratio measured at 10.5T vs 7T for these coils in the phantom is approximately equal to and slightly less than the intrinsic tSNR ratio measured with GRE EPI in the same phantom and with the same coils. This is expected only if temporal instabilities in the instrument including the RF coil are negligible and/or approximately equivalent. Thus, the 10.5T instrument has a significantly higher tSNR compared to the 7T scanner used (Siemens 7T Terra) as measured with GRE EPI and the acquisition parameters employed here. Therefore, performance of this 10.5T scanner is not a limiting factor in GRE BOLD fMRI studies. Should the tSNR ratio for 10.5T vs 7T *measured in the brain* be less than what is observed in the phantom, as is the case (see below), the cause cannot be ascribed to higher instrumental instabilities of the 10.5T system.

In order to avoid the complexity of calculating tSNR in a task-based paradigm, we acquired “resting state” data without a task or stimulus both at 10.5 and 7T, with identical acquisition parameters to those reported in Figure 2 in the manuscript and the phantom data (Figure S1.1). The RF coils employed were identical to the phantom data presented above. Figure S1.3A illustrates tSNR maps calculated for the 40 slices acquired at 10.5T (left column) and 7T (right column) in a coronal slab over the visual cortex with our off-line standard processing that included DORK^12^ for a ∼4 min run. Figures S1.3B and S1.3C illustrate an expanded view of 3 slices from these data for the two magnetic fields. Quantified over 9 such ∼4 min runs, tSNR ratio calculated in this one subject for all slices was 1.19 (tSNR=5.39±0.59 and 4.54±0.07 for 10.5T and 7T, respectively; errors reflect the variation among different runs). For the 3 slices selected in the expanded view (Figure S1.3B and S1.2C), the ratio was 1.5, 1.4 and 1.4 (given in the figure) although towards the surface of the brain tSNR became more comparable between the 10.5 and 7T datasets. For a second subject, the observed 10.5T vs 7T tSNR ratio was 1.23, reflecting tSNR of 5.50 ±0.33 and 4.47±0.14 for 10.5T and 7T respectively). Note that these numbers reflect tSNR without NORDIC denoising.

**Figure S1.3:**
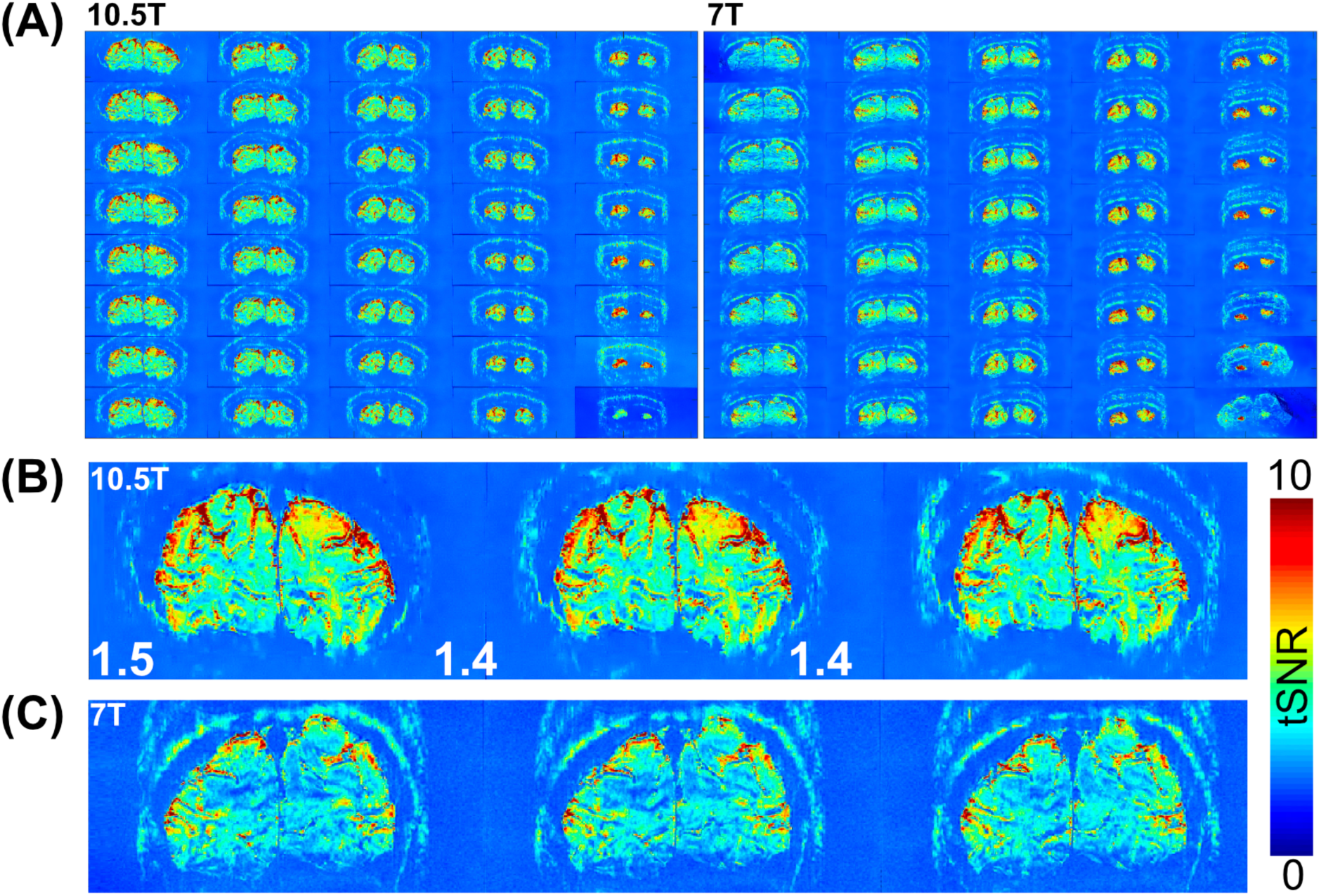
tSNR in coronal slices over a slab in the human visual cortex at 10.5T and 7T. (A) illustrates tSNR for all 40 slices at 10.5T (left column) and 7T (right column). Panels B and C show an expanded view of 3 slices from these data. Data are from the same subject. For the expanded view the slices were shifted by two to get a better match. The numbers at the bottom of panel B

Since the tSNR data are obtained with virtually the same TE of 24ms, the 10.5T suffers a larger T2* dependent signal loss than 7T. We corrected for this to get an “intrinsic” tSNR value, as was done in the phantom. An estimate of this can be obtained, for example, using gray matter T2* values measured in the visual cortex with 0.37 mm isotropic resolution (10.5T T2*=21.2 ms and 7T T2*=30.7ms, see manuscript Figure 1) which yields an increase of the tSNR ratio by factor 1.4 for the 10.5T/7T tSNR ratio in the brain putting the T2* corrected “intrinsic” tSNR ratio for the brain in the range ∼1.8. This is an approximate value since tSNR values we provide are slice averages and are not tissue specific. Note that the T2*-corrected tSNR ratio would also be the tSNR ratio that would be measured if the data were collected using the TE=T2* criteria for the two different fields, instead of using the same TE; in that case, at each field, there would be signal loss by factor *e*^−1^, which would cancel out in the ratio.

The upper bound for the T2* corrected “intrinsic” tSNR ratio is 2.75 (i.e. what was measured for this parameter in the phantom), which would be reached if thermal noise is fully dominant. However, the numbers obtained in the brain are markedly less than this upper bound. Therefore, the data on tSNR ratios suggest that thermal noise is substantial but not necessarily dominant in the brain at the spatiotemporal resolutions used in these acquisitions. In agreement with this, NORDIC denoising increased the slice averaged tSNR by ∼2 fold at 10.5T and somewhat more at 7T for the two subjects. NORDIC suppresses (reduces) zero-mean Gaussian noise that includes thermal noise. Hence the data indicate that zero-mean Gaussian noise constitutes ∼50% of the “noise” of the fMRI time series in the brain under these acquisition conditions. This, of course, is not surprising given the fact that the spontaneous oscillations of neuronal origin associated with resting state networks are a source of temporal fluctuations of signal amplitude in an fMRI time series^6,13,14^. In addition, although DORK does correct for respiration induced oscillations, it does not necessarily do so fully (e.g.^15^) and does not necessarily impact all other sources of non-neuronal structured noise. There are, however, a host of other approaches that can be brought to bear on reducing such non-neuronal sources of signal fluctuations (e.g.^2,4,7,8^ and references therein) and this remains to be explored fully both for 10.5T and 7T. A corollary of this conclusion is that even at this high resolution, it would be possible to detect the spontaneous oscillations that form the basis of resting state networks.

We note that the above numbers are based on averages over the slice while tSNR is not uniform through the slice and displays structure reminiscent of cortical anatomy. However, a detailed study with tissue segmentation etc. is beyond the scope of this paper and will be covered in the future.

### 6.2. Impact of RF Coils on Functional maps

This was an issue raised in the review process and as such we provide a detailed explanation below how this was dealt with. When evaluating fMRI maps (*not* just Δ*S*/*S*) at two different magnetic fields, we must consider coil performance because of the impact on SNR, and hence on fCNR. However, how does one compare coils built for two different magnetic field strengths? One cannot just focus on the number of channels, for example, or the coil layout. RF coils with the same number of channels and the same coil layout can and do perform differently at two different magnetic fields (e.g. ^10^). However, they can be compared using the “SNR Performance” metric which defines the SNR captured by the coil relative to the theoretically maximal SNR available at a given magnetic field (i.e. ultimate intrinsic SNR (uiSNR)). The coils used at the two different magnetic fields may have totally different configurations; but if they have the same “SNR Performance” then they capture the same fraction of the uiSNR available at the different magnetic fields.

In this respect, the coils employed in functional images (i.e., unless otherwise specified, CMRR built 10.5T 80-channel receive and CMRR built 7T 64-channel receive arrays) had approximately the same “SNR Performance” ^10^ (see also Figure S2). These coils both had a head-conformal, 64-channel inner receive-only array of identical coil layout but tuned to the different frequencies. However, the 10.5T coil used the 16-channel transmitter as a receiver as well, bringing up the receive channel count to 80. Despite the different number of receive channels, however, the fraction of the uiSNR captured by these coils operating at their respective frequencies were approximately the same (Figure S2). Of course, the uiSNR is much higher at 10.5T compared to 7T but the “SNR Performance”, i.e. the efficiency with which that uiSNR was captured, was the same for these two coils, making them ideal for this 7T vs 10.5T comparison for fMRI. If anything, the 64-channel array at 7T performed slightly better towards the center than the 10.5T RF coil in terms of SNR efficiency and conferred an advantage to 7T compared to 10.5T.

**Figure S2:**
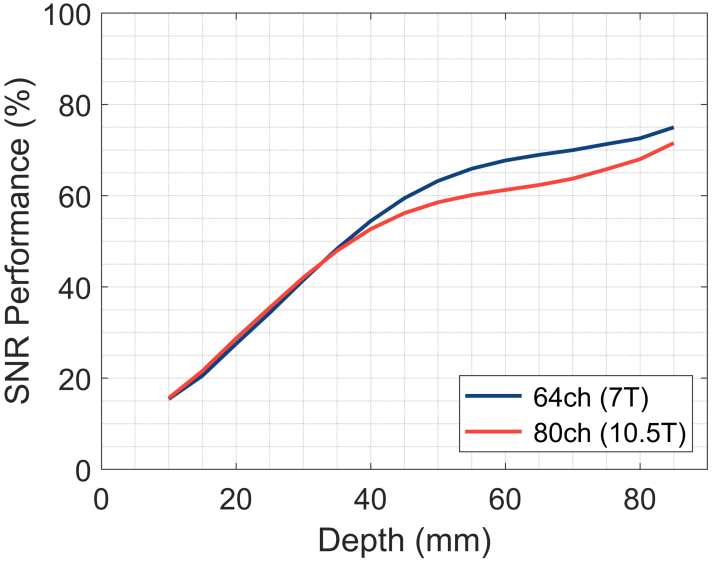
SNR performance (Array SNR divided by uiSNR, reported as %) of the CMRR built 7T 64-channel receive coil *versus* the 10.5T 80-channel receive coil employed in these studies. SNR performance is shown as a function of depth averaged in a function of depth averaged in concentric shells. (see references 11,12 for details). The 7T 64 channel receive array was built specifically to duplicate the coil layout of the inner 64 receive only array of the 10.5T 80 channel coil.

Functional images at **both magnetic fields** can improve with better RF coils, especially in the peripheral regions where the uiSNR is intrinsically very high but RF coils capture a small fraction of it. 7 Tesla has had a long history of RF coil development and optimization, while 10.5 Tesla is new and the coils available are the first generation. Therefore, the room for improvement is higher at 10.5T.

### 6.3. Percent BOLD signal change versus RF receive and/or transmit coil performance

BOLD effect, evaluated as the signal increases (Δ*S*) induced by altered brain activity as a ratio of the steady-state image signal intensity *S* in the fMRI time series (i.e. = (Δ*S*/*S*)), is strongly magnetic field dependent but is **independent** of instrumentation details, such as RF coils or gradients. Δ*S*/*S* is given by the product of TE and the 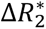, where 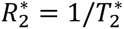 and 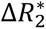 represents the change induced in 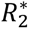 by the altered neuronal activity. Although this derivation was given before^16^, we reproduce it here in greater detail given questions raised in the review process about the role of the receive and transmit RF coils on this parameter.

The BOLD effect reflects a change in the 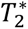 of individual voxels^16^ coupled to neuronal activity. Explicitly including the RF coil sensitivity profile 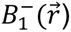, where 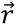 designates spatial coordinates over the sample, one can write

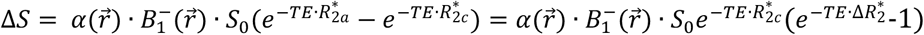

where *TE* is the delay time, 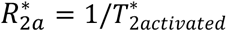 and 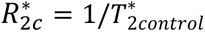 where “activated” designates the task/stimulus “on” state, “control” is the state to which the task/stimulus state is referenced, and 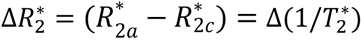. *α* is a constant for the experiment, determined as a function of T1, TR and the flip angle for that voxel; thus, *α* depends on the transmit coil used and when transmit *B_1_* (*i*. *e*. 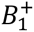) is spatially nonuniform, *α* becomes a function of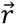 as well. *S*_0_ is the thermal equilibrium magnetization. The expression 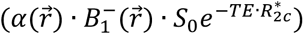 is the steady state signal intensity *S* detected for the “control” condition in the fMRI time series in the absence of a task or stimulus. Since task/stimulus induced changes in fMRI are very small, i.e. 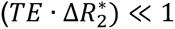, the equation above reduces to

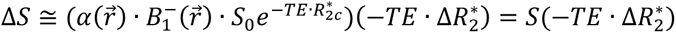

leading to 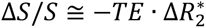

Note that in this expression 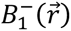 as well 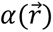 and hence 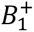, drop out of the equation for Δ*S*/*S*. Instead, Δ*S*/*S* is only proportional to *TE* and 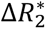, as previously shown^16^; the latter is a fundamental property of the voxel and cannot depend on the RF coil transmit or receive properties or the details of any other hardware employed. BOLD modeling aims to calculate this 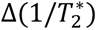 by considering the effect of the field inhomogeneities around blood vessels, vascular anatomy, complex set of parameters such as hematocrit, oxygenation level in arterial and venous blood vessels, blood flow and oxygen consumption rate, and changes in these parameters mediated through neurovascular coupling.

Although it is conceptually evident, as shown above, that Δ*S*/*S* must be independent of RF coils, we experimentally demonstrate that as well in this paper. In Figure 2 of the manuscript, comparing Δ*S*/*S* at 7 vs. 10.5 Tesla, subject S3 data used a different 7T coil (64 channel receive 16 channel transmit (64Rx/16Tx) acquired from MR CoilTech Ltd., UK), while the other two subjects employed a CMRR built 64channel receive, 1-channel transmit RF coil, which was built specifically to replicate the design of the 10.5T coil employed in this study. The data demonstrate experimentally that the parameter Δ*S*/*S*, as expected, did not change.

The two different 7T coils employed are compared for “SNR Performance” defined as the fraction of the theoretically maximal SNR (i.e. ultimate intrinsic SNR (uiSNR)) captured by the coil (Figure S3). This figure also shows the same data for the industry standard, 32-channel receive,1 channel transmit 7T coil supplied by NOVA Medical (Wilmington MA, USA). The data in Figure S3 were obtained using a phantom mimicking human head electrical properties and dimensions (phantom described in detail in^9,10^). Because of the symmetry involved, SNR performance was averaged in 1 cm thick concentric shells, which at the periphery conform to the phantom’s outer contour and is then progressively shrunk to obtain a 1-dimensional plot, as a function of depth from the phantom surface to the phantom middle (see also ^9,10^). This figure clearly illustrates that the MR CoilTech 64 channel 7T coil has superior “SNR Performance” compared to the CMRR 64 channel 7T coil, and also incidentally the NOVA 32 channel coil as well. In other words, all else being equal images obtained with the MR CoilTech coil will have higher SNR than the other 2 coils, as much as 30% more in the periphery and 13% more in the center relative to the CMRR 64channel coil. Nevertheless, new data on Δ*S*/*S* measured with the CoilTech and CMRR 64 channel coils is the same, yielding ∼2-fold lower PSC at 7T relative to the 10.5T. Note that the CMRR 7T coil was not built to optimize performance at 7T but rather duplicated the design choices that were made for building the first generation 10.5T coil, which was employed in this paper; the design choices propagated from 10.5T included a self-decoupling coil circuitry which does not work well at 7T for coil sizes that correspond to receive coils.

**Figure S3:**
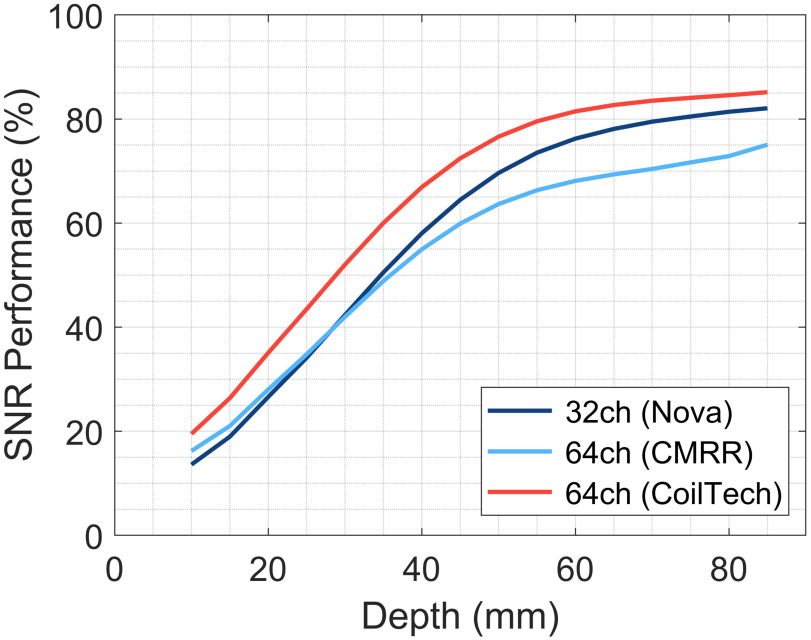
7 Tesla SNR performance (defined as Array SNR divided by uiSNR) reported as % for 3 different 7T coils, as a function of depth as in Figure S2. CoilTech designates a 64-channel receive/16 channel transmit coil from MR CoilTech Ltd. (UK)

### 6.4. High-Performance Gradients in fMRI

The present data were obtained without high-performance head gradients. Such a gradient (i.e. the Impulse gradient from Siemens) is scheduled to be installed at 10.5T in mid 2027. Because of questions raised about the possible implications of high-performance gradients, we include a discussion here.

High-performance gradients do *not* impact the BOLD mechanism and hence the BOLD effect quantified as Δ*S*/*S* (i.e. percent signal change or PSC) and reported in this paper. They do impact fMRI experiments, but this impact is complex and nuanced unlike, for example, their effect on diffusion weighted MRI (dMRI) where they provide a major improvement in SNR.

The SNR advantage of high-performance gradients in dMRI arises because they enable a substantial reduction in TE, the delay inserted after signal excitation and prior to the start of image acquisition where diffusion encoding takes place. During this delay, SNR is lost exponentially with TE and a shorter TE means higher SNR. As such, the motivation for the development of high-performance gradients came primarily from dMRI applications starting with the Human Connectome Project ^17,18^.

fMRI also has a TE, a delay prior to image acquisition used for encoding BOLD contrast (*gradients are not employed during this period)*. While in dMRI, one strives to make the TE as short as possible, in fMRI TE is set ≅T2* with the goal of optimizing fCNR ^16^. Thus, in BOLD fMRI, optimum TE is determined by the T2*. *As long as the acquisition achieves approximately this optimum TE*, high-performance gradients do not provide an SNR gain through TE, as they do in dMRI.

The primary gain of the high-performance gradients in fMRI is to reduce **blurring** and only **in one dimension**, the phase encode dimension, as demonstrated previously^19^. The high maximal gradient strength (Gmax) enabled by high-performance gradients shortens the duration of the echo train length (ETL) in EPI imaging (hence reduces blurring). However, the high Gmax means the image acquisition uses a higher bandwidth (*BW*), which reduces SNR since SNR depends inversely on the 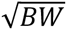. Thus, in fMRI, to a *first approximation*, high-performance gradients do not improve SNR (may in fact reduce it) and hence do not improve fCNR, compared to a “standard” gradient, provided that the TE≅T2* condition is met (a condition that is fulfilled in this paper) and all the echoes collected in the EPI echo train have sufficient SNR. The latter caveat leads to blurring if the later echoes in the ETL with the standard gradient do not have good SNR and thus fail to capture high-frequency information in k-space needed for high-resolutions. The severity of this will depend on the magnetic field strength and spatial resolutions sought; while at the higher magnetic field signal decay during the ETL is faster, the inherently higher SNR can enable even the later echoes to have sufficient SNR to capture the high-frequency information and preclude extensive blurring. We in fact provide data that the blurring in our 0.35 mm isotropic data is minimal even without the high-performance gradients (Figure 4d) unlike what was reported previously at 7T where blurring became substantial starting at 0.6 mm isotropic resolution with the same “body-gradient” performance employed in the 10.5T studies reported in this paper. Of course, pushing to even higher resolutions, however, will need high-performance gradients even at 10.5T.

All the benefits of high-performance gradients in fMRI attained through the reduction of the ETL in EPI can also be achieved using higher image acceleration through parallel imaging. Acceleration with parallel imaging reduces the ETL and suffers an SNR loss equivalent to that coming from the higher bandwidth used with high-performance gradients to achieve shorter ETL. In other words, if you reduce ETL in EPI by factor of, say 2, either using high-performance gradients or 2-fold acceleration through parallel imaging, SNR will decrease by 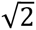. In case of parallel imaging, there will be an additional degradation by the g factor^20^. With respect to parallel imaging, however, higher magnetic fields provide a substantial advantage because they provide significantly higher acceleration capability^9,21,22^ while keeping g∼1. Thus, one can accelerate more at 10.5T compared to 7T for a given g-factor, as we demonstrated previously ^9,10^. In these studies, we have used a factor of 4 in-plane acceleration (iPAT in Siemens terminology) while typically at 7T, including for example in the paper^19^ reporting on the impact of high-performance gradients on fMRI, one uses 3 or less in-plane acceleration.

**Figure S5:**
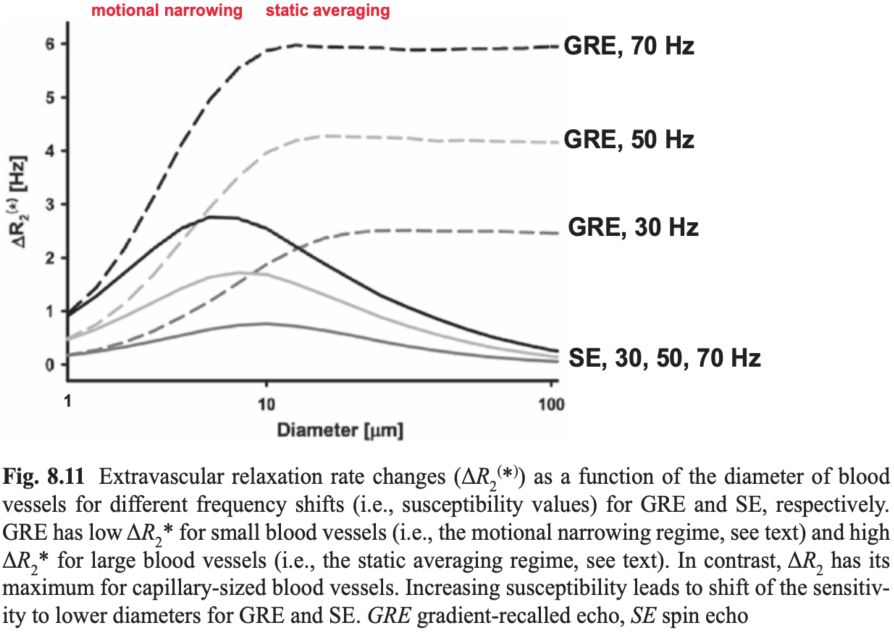
the GRE and SE BOLD effect as a function of diameter with infinite cylinders used as approximating blood vessels. The figure and its caption are directly reproduced from reference 23.

Ideally, of course, it is best to have high-performance gradients at the very high magnetic fields, adding to the high acceleration potential through parallel imaging for improved performance. This is in process for 10.5T.

### 6.5. Spin Echo vs Gradient Echo BOLD

For convenience, Figure 8.11 from reference ^23^ is reproduced in Figure S5, demonstrating that SE BOLD is inherently smaller than GRE BOLD for a given vessel size, because of diffusion averaging of the BOLD effect. Similar results were shown by others as well (e.g. ^24,25^)

### 6.6. The resolution effect on large vein contributions to GRE BOLD fMRI

If one considers an infinite cylinder as an approximation for a blood vessel with magnetic susceptibility difference Δ*χ* relative to its surrounding, the applied magnetic field of *B*_0_ will be perturbed and deviate from it outside the cylinder.^25–27^ This perturbation, expressed as 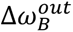 in rad/s, relative to the magnetic field distant from the cylinder, *ω*_0_, will be given by the equation

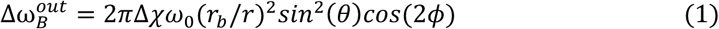

In the above equations, *ω*_0_ is *B*_0_ expressed as Larmor frequency in rad/s (i.e. *ω*_0_ = *γB*_0_ where *γ* is the gyromagnetic ratio), *θ* is the angle between the applied magnetic field *B*_0_ direction and the cylinder axis in a plane that contains them both, *r_b_* is the cylinder radius, *r* is the distance between the point of interest and the center of the cylinder cross-section in the plane normal to the cylinder, and *φ* is the angle between this vector 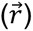 and the component of B_0_ in this plane (Figure S6.1).

**Figure S6.1:**
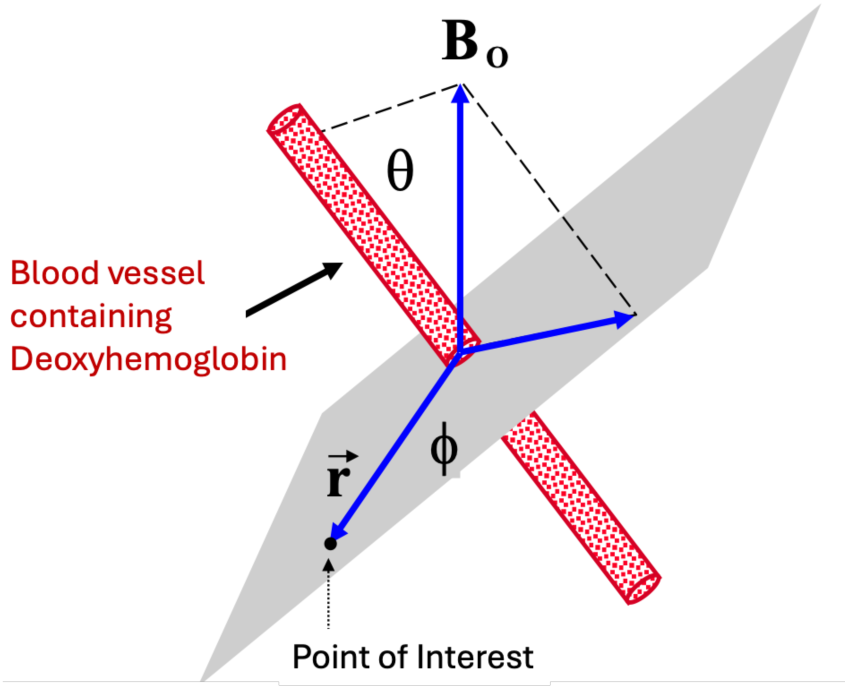
Definition of the variables given in Eq.

The extravascular BOLD effect originates from the intra-voxel inhomogeneity caused by 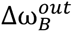 outside the deoxyhemoglobin-containing blood vessels. BOLD fMRI signals arise when this intra-voxel inhomogeneity is altered due to small changes in Δ*χ* and/or *r_b_* induced by alterations in neuronal activity. Therefore, the mechanistically critical phenomenon in extravascular BOLD effect is the intra-voxel inhomogeneity caused by the inhomogeneous 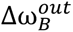.

Lumping all the terms in Eq. 1 other than (*r_b_*/*r*) into a function designated as *K*(*θ*, *φ*, Δ*χ*), one can write

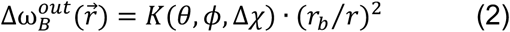

If we consider the problem only in one-dimension along the vector 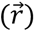, limiting ourselves for convenience to a one-dimensional imaging so to speak, the gradient of the magnetic field with respect to *r* will be given by the derivative of Eq. 2,

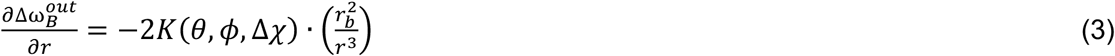

Across a one-dimensional voxel of dimension *δr* then, for small *δr*, the magnetic field variation across the voxel (i.e. intra-voxel magnetic field inhomogeneity) will be given by:

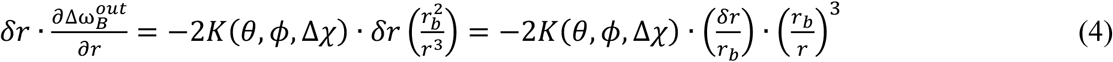

This intra-voxel magnetic field variation will induce intravoxel-dephasing during a delay TE in the GRE BOLD fMRI acquisition, reducing the amplitude of the signal detected for this voxel. This is the BOLD effect. Alterations in neuronal activity will induce changes in the intra-voxel dephasing and hence in the signal amplitude of the voxel, giving rise to the functional signals in fMRI.

The variation of 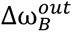 within a voxel of dimension *δr* at the surface of the blood vessel where *r* = *r_b_* will be

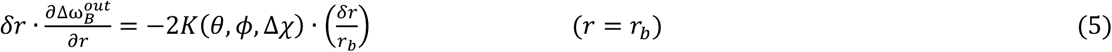

Thus, the gradient of the magnetic field outside the blood vessel is steepest at/near the luminal boundaries of the blood vessel where *r* = *r_b_*, and will decrease from that value as the cube power of (*r_b_*⁄*r*) with increasing distance away from the blood vessel. As the resolution gets higher (i.e. *δr* gets smaller) relative to the blood vessel radius, in the limit (*δr*⁄*r_b_*) ≪ 1, the intra-voxel variation of the magnetic field for all voxels outside the cylinder will become negligible. Hence the signal amplitude decreases during TE (i.e. the extravascular BOLD effect manifested as an amplitude modulation) will also migrate beyond detection limits. Note that extravascular perturbations of the magnetic field outside the deoxyhemoglobin-containing blood vessel given by Eq.1 still exist and can be detected as a phase variation over spatial coordinates but will not be detected in a standard GRE BOLD fMRI experiment which looks at signal amplitude changes.

For blood vessels that do not fully satisfy this condition (e.g. *δr*∼*r_b_*), the BOLD effect will persist near the blood vessel boundary where the gradient is large. However, with increasing separation from the blood vessel, the intra-voxel magnetic field variation, given by Eq. 4, will rapidly decrease according to (*r_b_*⁄*r*)^3^, vanishing when 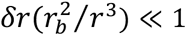. In other words, as (*δd*/*r_b_*) gets smaller, the extravascular BOLD effect (detected as signal amplitude associated with a voxel), starts progressively shrinking towards the blood vessel and appears in close proximity to the blood vessel boundary where the gradient is steepest. This is what is observed in Figure 4b (right-most panel labeled S4 expanded) of the paper.

The condition (*δr*⁄*r_b_*) ≲ 1 can be satisfied for different size draining veins at submillimeter spatial resolutions. But it will be far from being satisfied for capillaries (diameter^28^ ∼5.1±0.84 µm) and postcapillary venules (few tens of microns in diameter). Hence, as the spatial resolution increases, the microvascular BOLD effect remains unaffected while the large vein contributions decrease, leading to increased domination of the microvascular contribution at the very high magnetic fields.

**Figure S6.2:**
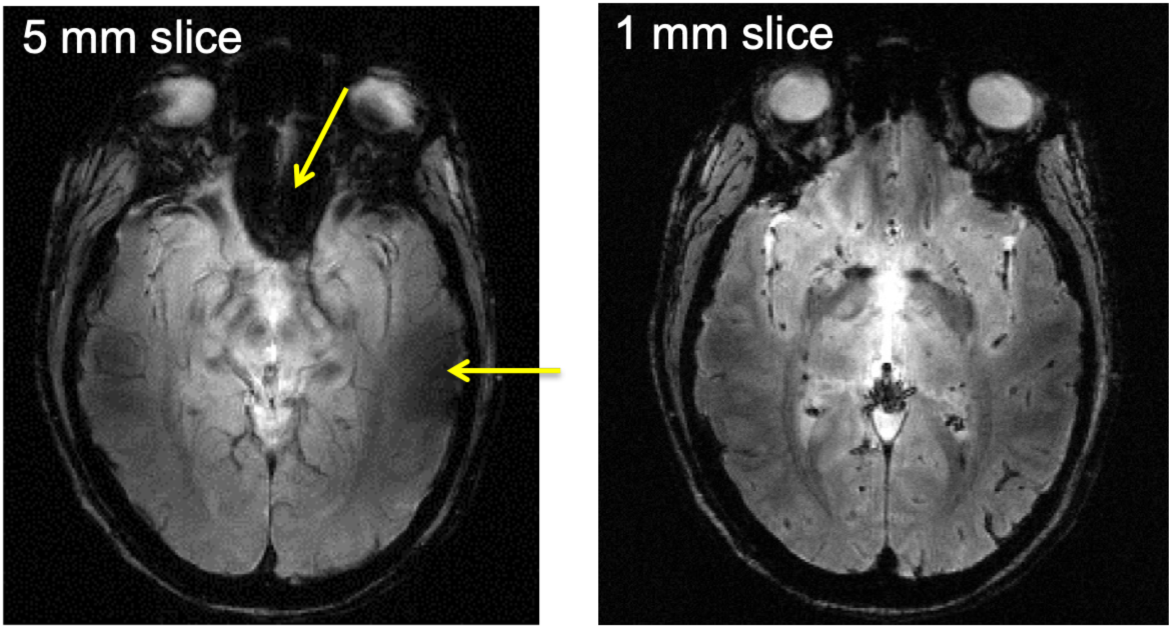
Gradient recalled echo images in the human brain of a slice through a region near nasal cavities and sinuses. The left panel has a lower resolution with a 5 mm slice thickness. Total signal dropout is observed in the frontal areas and in addition, signal intensity reduction (hence darker image intensity) is observed near the ears (yellow arrows). These are well known effects associated with the presence of magnetic field gradients near the air-filled cavities in the human head. Reducing the voxel volume by cutting down the slice thickness eliminates the signal dropout in the frontal areas and improves image intensity near the ears

There is of course also an intravascular BOLD effect. However, the T2 of deoxyhemoglobin-containing blood decreases rapidly with increasing magnetic fields so that at fields ≳7 Tesla, at TEs that approximate the tissue T2, as employed in GRE BOLD fMRI, the blood signal disappears^29^. At lower magnetic fields, the intravascular BOLD effect remains significant (e.g. at 3 Tesla it was estimated to be ∼50% of the BOLD signal^30^) and can be a source of draining vein signals. Thus, the suppression of draining veins through high resolutions requires the high magnetic fields because of the high SNR needs of high-resolution imaging but also for eliminating the intravascular BOLD effect.

The resolution effect discussed above can also be conceptualized using simple GRE images of a brain slice where signal intensity dropouts occur near the air-filled cavities in the frontal regions because of magnetic field inhomogeneities. Analogous to the EV BOLD effect, this intensity dropout is induced by the susceptibility difference between brain tissue and air. It is well recognized that at high enough resolutions, as the voxel dimensions get smaller compared to the distances over which magnetic fields vary due to this susceptibility difference, the intra-voxel signal variation becomes smaller, and the signal dropouts disappear. This is illustrated in Figure S6.2 with an image acquired with 5 mm slice thickness versus a 1 mm slice thickness.

### 6.7 7T 0.8 mm activation maps in native (non-upsampled) resolution

**SUPPLEMENTARY FIGURE S7:**
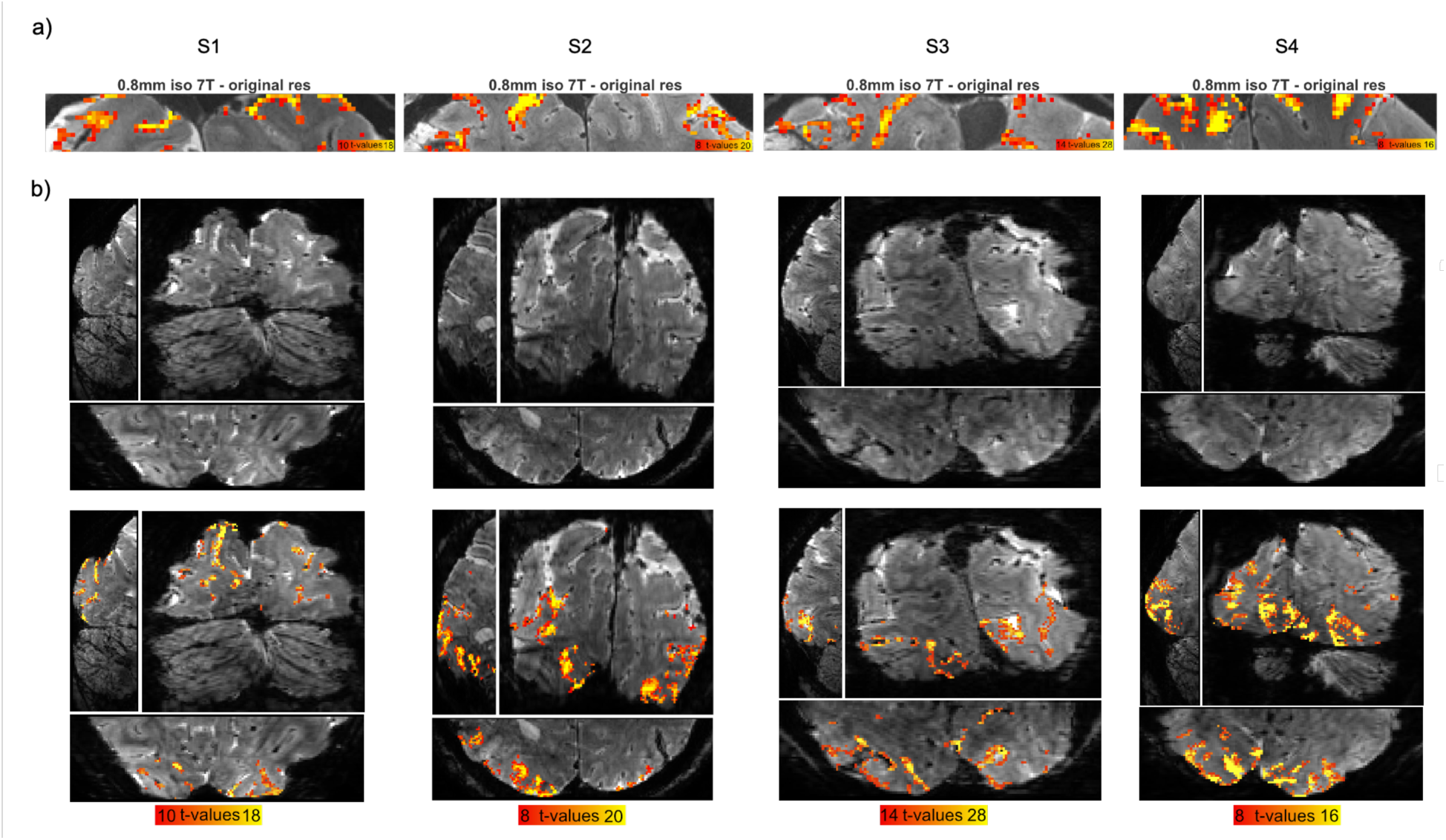
a) 7T 0.8 mm activation maps from Figure 4c shown here in native (non-upsampled) resolution for reference. **b)** Additional example slices (sagittal, coronal and axial orientations) of 0.8 mm activation maps (cluster threshold of 10) in their native resolution (lower panel). The underlay is the mean 0.8 mm EPI data averaged across the time series of one run, which is shown without activation for reference in the top panel.

### 6.8. 0.35 vs 0.8 mm iso functional images: comparing ramping profiles across cortical depths

As mentioned in the main section, to estimate the slope of PSC responses ramping towards the pial surface in GRE BOLD fMRI, we fitted linear models to single block PSC responses across depths (see Methods). The average slope of the ramping profile displaying increasing PSC towards the pial surface (i.e. towards CSF) – typical of GRE BOLD fMRI – was significantly shallower for the 0.35mm 10.5T fMRI data compared to the 0.8 mm 7T fMRI images, as evaluated with two-sample t-tests (S1: (mean±Std across trials) 0.35 mm: 3.85±1.76, 0.8 mm: 5.93±0.97, t(22)=-3.61, p=0.002; S2: 0.35 mm: 6.01±1.24, 0.8 mm: 7.47±2.03, t(22)=-2.12, p=0.045; S3: 0.35 mm: 8.12±2.09, 0.8 mm: 12.63±2.87, t(22)=-4.41, p<0.001; S4: 0.35 mm: 1.81±5.92, 0.8 mm: 5.75±2.06, t(22)=-2.18, p=0.040 – Figure S8). This was quantified by fitting a line to the PSC laminar profile elicited by each block and estimating its slope (see Methods). This result was primarily driven both by the apparent increase in evoked PSC amplitudes in the inner depths and the decreased amplitudes towards the pial surface.

**SUPPLEMENTARY FIGURE S8:**
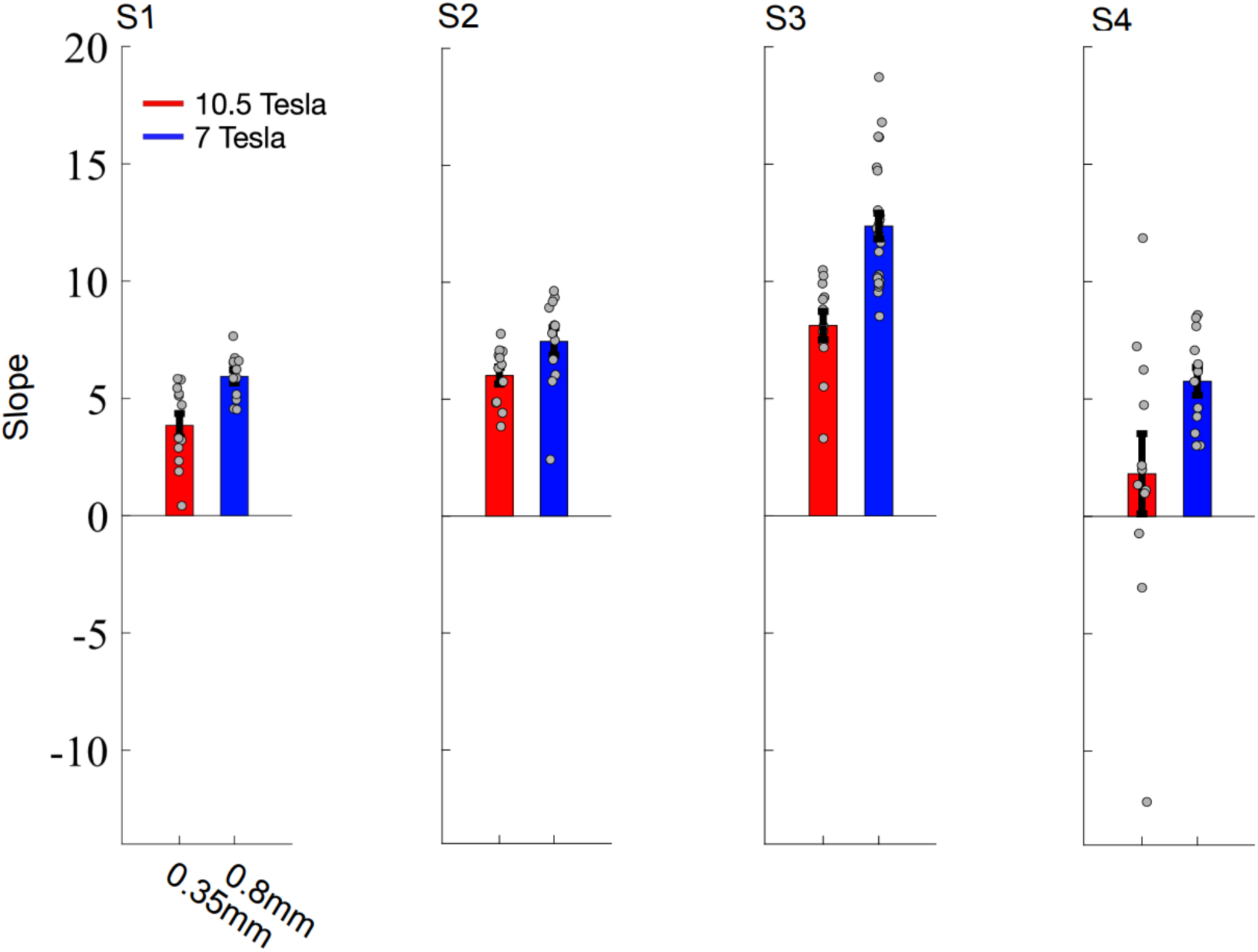
the bar plots depict the average slopes estimated across single blocks (grey dots) to quantify the ramping profiles across cortical depths for the 0.8 mm iso (blue) and 0.35 mm iso (red) images (see Methods).

### 6.9. Zoomed versions of Figure 4

**SUPPLEMENTARY FIGURE S9.1:**
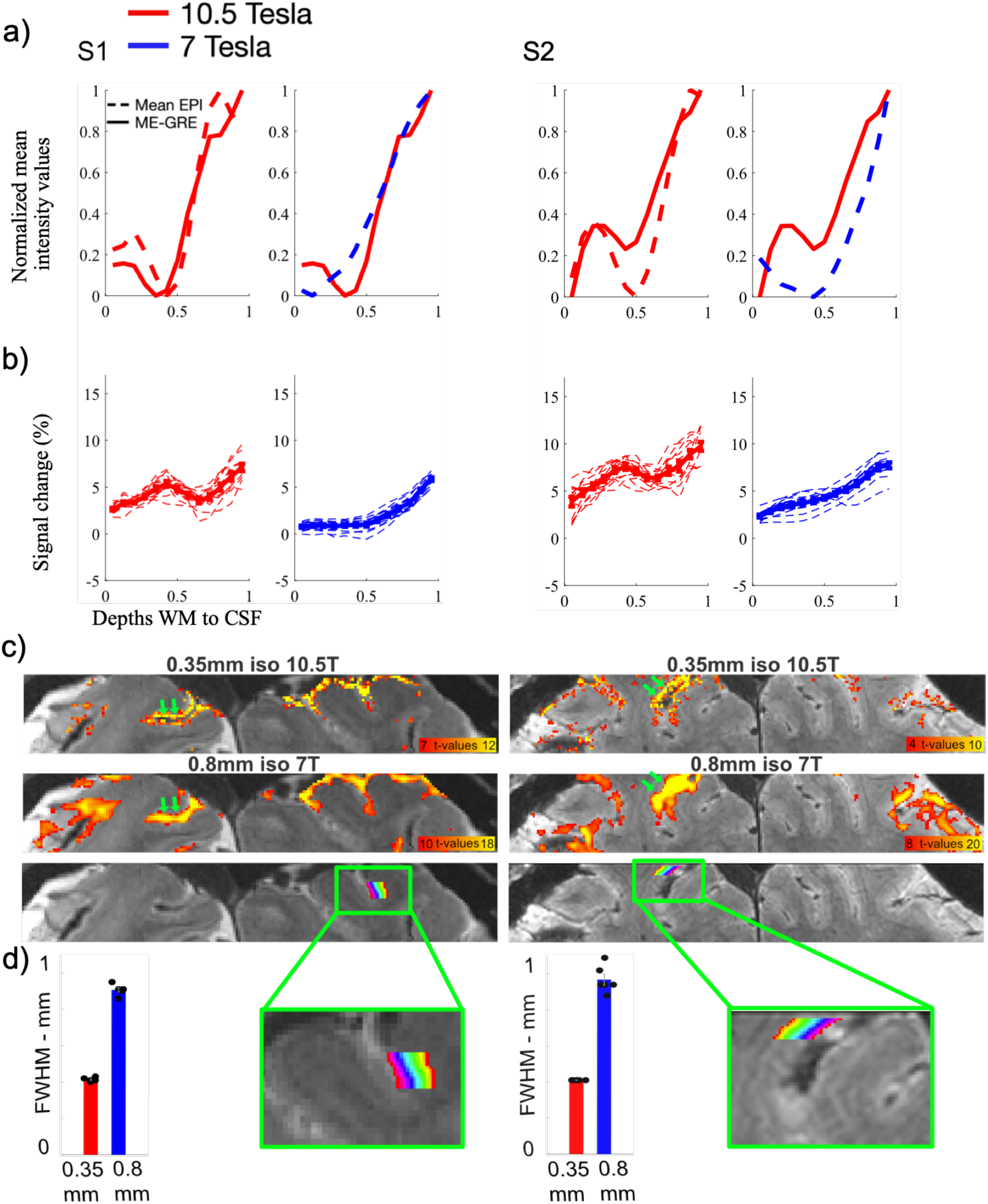
Data from the first two subjects (S1 and S2) from Figure 4 shown here as a zoomed version. **Panel (a)** depicts laminar profiles of normalized (min=0, max=1) signal intensities for the T2*-weighted acquisitions (mean EPI and MEGRE images). The dashed lines refer to the mean EPI intensity profiles (red: 10.5T 0.35 mm; blue: 7T 0.8 mm). The MEGRE data are from 10.5T and have 0.37 mm isotropic resolution (solid red line, included in both panels of each subject). The dip in the middle depths in the 10.5T MEGRE and 0.35 mm EPI profiles identifies the location of the Stria of Gennari, which is not visible in the corresponding 0.8 mm 7T mean-EPI profiles. **Panel (b)** shows the functional PSC laminar profiles for the target condition. For both subjects, the 0.35 mm PSC profiles (in red) demonstrate a bump co-localized with the dip seen in the mean-EPI and MEGRE anatomical traces. **Panel (c)** The top and middle rows show activation (T-maps, cluster threshold of 10) for the 0.35 and 0.8 mm acquisitions, respectively. The green arrows point to example locations where activation across adjacent sulcal banks is separable at 0.35 mm but not 0.8 mm resolution. Bottom row illustrates cortical layers used to extract laminar profiles in panel (a) from an example slice in each subject (d) Bar plots

**SUPPLEMENTARY FIGURE S9.2:**
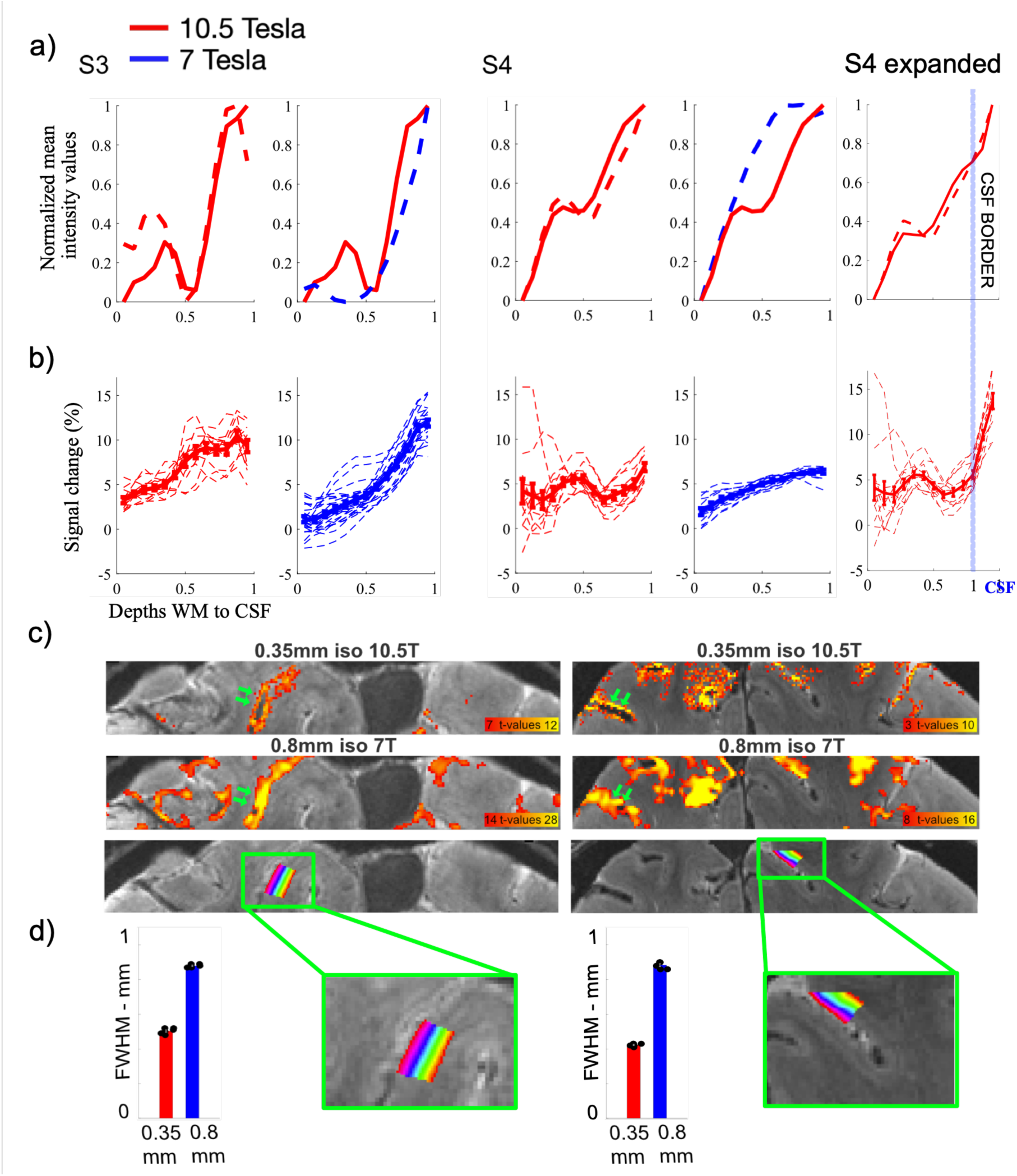
As in Figure S9.1 above, but for subjects S3 and S4. In addition, this figure contains a fifth column (rightmost column) in **Panels (a)** and **(b),** labeled **“S4 expanded”**. This panel shows the PSC in the 10.5T 0.35 mm data, rapidly increasing immediately beyond the boundary between cortex and CSF (light-blue vertical line).

